# Cis-delivering releasable IL-15 superagonist enhances antitumor immunity in cold tumors by invigorating preexisting CD25^+^CD8^+^ T cells

**DOI:** 10.1101/2025.08.14.670432

**Authors:** Qiongya Zeng, Wencheng Kong, Ying Qin, Luyao Song, Nan Liu, Zexin Liu, Chenghao Ji, Gen Qi, Wenqiang Shi, Huili Lu

## Abstract

Antibody-mediated cis-delivery and trans-delivery both direct cytokines to tumors and have been extensively investigated in clinical trials. However, a comparative analysis of their differential effects on cytokine activity is still lacking. In this study, we initially verify that cis-delivery of cytokine demonstrates a markedly stronger antitumor effect than trans-delivery, but it also exhibits certain drawbacks, including severe toxicity, insufficient activation of CD25^+^CD8^+^ T cells, and enhanced stimulation of intratumoral regulatory T cells (Tregs). To further address these issues, we developed a conditionally releasable and cis-delivering IL-15 immunocytokine (termed PMIS), which can selectively release a free IL-15 superagonist within tumors rather than immobilizing IL-15 to PD-1^+^ Tregs, thereby potently stimulating CD25^+^CD8^+^ T cells. Mice treated with PMIS showed significantly reduced systemic toxicities while achieving notably stronger antitumor effects. Administered either alone or in combination with other therapies, PMIS exhibits great potential for inhibiting orthotopic cold tumor and its metastases. Mechanistically, the significant activation of the pre-existing intratumoral CD25^+^CD8^+^ T cells and the improved CD25^+^CD8/Treg ratio contribute to the enhanced antitumor response of PMIS. These findings underscore the indispensable role of CD25^+^CD8^+^ T cells in cis-delivering IL-15 and provide a promising strategy for overcoming resistance to therapies and effectively controlling advanced cold tumors.

## Introduction

Antibody-cytokine fusion proteins, also known as immunocytokines, have drawn growing attention for their unique mechanisms of action and notable antitumor effects, steadily establishing themselves as a rising star in the field of cancer immunotherapy(1). This fusion strategy not only extends the half-life of cytokines and reduce off-target toxicities but also leverages the combined strengths of both antibody and cytokine components to achieve synergistic antitumor effects(2). Early immunocytokines (currently under phase 2/3 trials) primarily target tumor-associated antigens, such as fibronectin, fibroblast activation protein (FAP), and epidermal growth factor receptor (EGFR)(3). Of note, immunocytokines targeting programmed death 1/programmed death-ligand 1 (PD-1/PD-L1) hold significant potential to overcome resistance to immune checkpoint blockade, thereby emerging as a key focus for development in recent years(4). For immunocytokines like anti-FAP/IL-2, anti-PD-L1/IL-15, and anti-EGFR/IL-10, the antibody components bind to tumor cells, while the cytokines bind to immune cells(5–7). This method of cytokine delivery is referred to as trans-delivery/presentation. In contrast, anti-PD-1-based immunocytokines employ a process known as cis-delivery/presentation, where both the antibody and cytokine target the same cells (T cell) (8).

Studies have demonstrated that anti-PD-1 could effectively cis-deliver cytokines to PD-1^+^CD8^+^ T cells, resulting in a more potent reinvigoration of intratumoral exhausted CD8^+^ T cells, thereby endowing anti-PD-1-based immunocytokines with superior antitumor potential(9, 10). Several anti-PD-1-based immunocytokines are currently under clinical investigation, demonstrating encouraging efficacy in patients with refractory advanced solid tumors (NCT05396391; NCT05460767; NCT04303858). However, the cis-delivery of cytokines still faces two significant challenges: 1) there remain considerable dose-limiting toxicities similar to those of the parental cytokines(11); 2) both intratumoral CD8^+^ T cells and regulatory T cells (Tregs) express high levels of PD-1, posing a risk of simultaneously stimulating Tregs while activating CD8^+^ T cells(12). The increased immunosuppressive effects of Tregs could negatively impact the antitumor activity.

Regarding safety issues, two primary strategies are under investigation: directly reducing cytokine activity, or employing a prodrug strategy to mask cytokine activity in normal tissues while selectively activating cytokine signaling within the tumor microenvironment (TME) (13). However, reducing the potency of cytokines often diminishes their antitumor effects, making it challenging to identify the optimal level of activity reduction that balances peripheral safety with effectiveness in tumor tissues. In prodrug strategies, a common approach to engineer a cytokine prodrug involves fusing a cognate receptor or high-affinity antibody fragment and inserting a tumor-associated protease substrate linker. This design enables the cytokine to restore its activity upon cleavage by the protease within the TME(14–16). It is important to note that, in this strategy, the cytokine remains fused with the anti-PD-1 after cleavage, which can still present the cytokine to Tregs and stimulate them. Therefore, it is crucial to mitigate the adverse effects of Tregs on the antitumor activity of cytokines cis-delivery.

IL-2 and IL-15 are the most extensively studied payloads for PD-1 cis-delivery immunocytokines. For IL-2, to reduce the stimulatory activity on CD4^+^CD25^+^ Tregs, various biased agonists of IL-2 have been developed. For example, Roche designed an IL-2 variant (IL-2v) with reduced binding to IL-2Rβγ and deleted binding to IL-2Rα (CD25). The PD-1/IL-2v immunocytokine effectively rescued effector T cells from Treg-induced suppression(17). In contrast, a recent study of Regeneron reported that IL-2Rα-biased binding can be beneficial and the engineered PD1-IL2Ra-IL2 construct stimulated more tumor-specific PD-1^+^CD25^+^ tumor-specific T cells than Tregs, leading to potent antitumor efficacy(18). However, the study did not further investigate the impact of intratumoral Treg stimulation on these effects. IL-15 is considered superior to IL-2 for its significantly weaker Treg-stimulating activity(19). Due to the anti-PD-1-mediated enrichment of cytokines to Tregs, it’s possible that a minor Treg-stimulating activity of a cytokine could be greatly enhanced. Therefore, it is critical to clarify whether the anti-PD-1/IL-15 immunocytokine would result in Treg stimulation in the TME and to explore viable solutions to address this concern.

In this study, we first compared the antitumor effects and safety profiles of cis-delivery versus trans-delivery of IL-15, demonstrating that cis-delivery did lead to enhanced Treg stimulation. To address this challenge, we innovatively masked IL-15 activity through steric hindrance to engineer a next-generation anti-PD-1/IL-15 prodrug. It can be specifically cleaved within the TME to release the anti-PD-1 and the free IL-15 superagonist. This strategy potently stimulates intratumoral CD25^+^CD8^+^ T cells, resulting in a higher CD25^+^CD8^+^/Treg ratio and enhanced antitumor effects with reduced toxicity. Overall, our work provides a new strategy and foundation for the development of next-generation anti-PD-1-based immunocytokines.

## Results

### Cis-delivery of cytokine shows superior antitumor efficacy to trans-delivery, yet this effect is compromised by Tregs

Both anti-PD-1-mediated cis-delivery and anti-PD-L1-mediated trans-delivery of cytokines demonstrate potent antitumor activities, however, there is currently no research directly comparing the differences in their antitumor potentials (Figure 1A). We first fused the IL-15Rα-sushi domain/IL-15 complex to the C-terminus of anti-PD-1 and anti-PD-L1 to construct αPD-1/IL-15 and αPD-L1/IL-15, respectively. In enzyme-linked immunosorbent assays (ELISAs), αPD-1/IL-15 demonstrated high-affinity binding to both human and mouse PD-1, comparable to that of the anti-PD-1 antibody (Figure S1A and S1B). The αPD-L1/IL-15 utilized in this study is one we reported previously(20), which exhibited a proliferative capacity in Mo7e cells similar to that of αPD-1/IL-15 (Figure S1C). We then evaluated the antitumor efficacy and safety of these two molecules in a mouse advanced tumor model. The results showed that 0.6 mg/kg of αPD-1/IL-15 exhibited a significantly stronger antitumor effect compared to 1 mg/kg of αPD-L1/IL-15 (Figure 1B), while 1 mg/kg of αPD-1/IL-15 demonstrated considerable toxicity, resulting in dramatic weight loss and eventually the death of all mice after two treatments (Figure 1C). These results indicate that anti-PD-1 mediated cis-delivery of IL-15 exhibits superior antitumor activity, but it is accompanied by severe toxicity.

**Figure 1.**
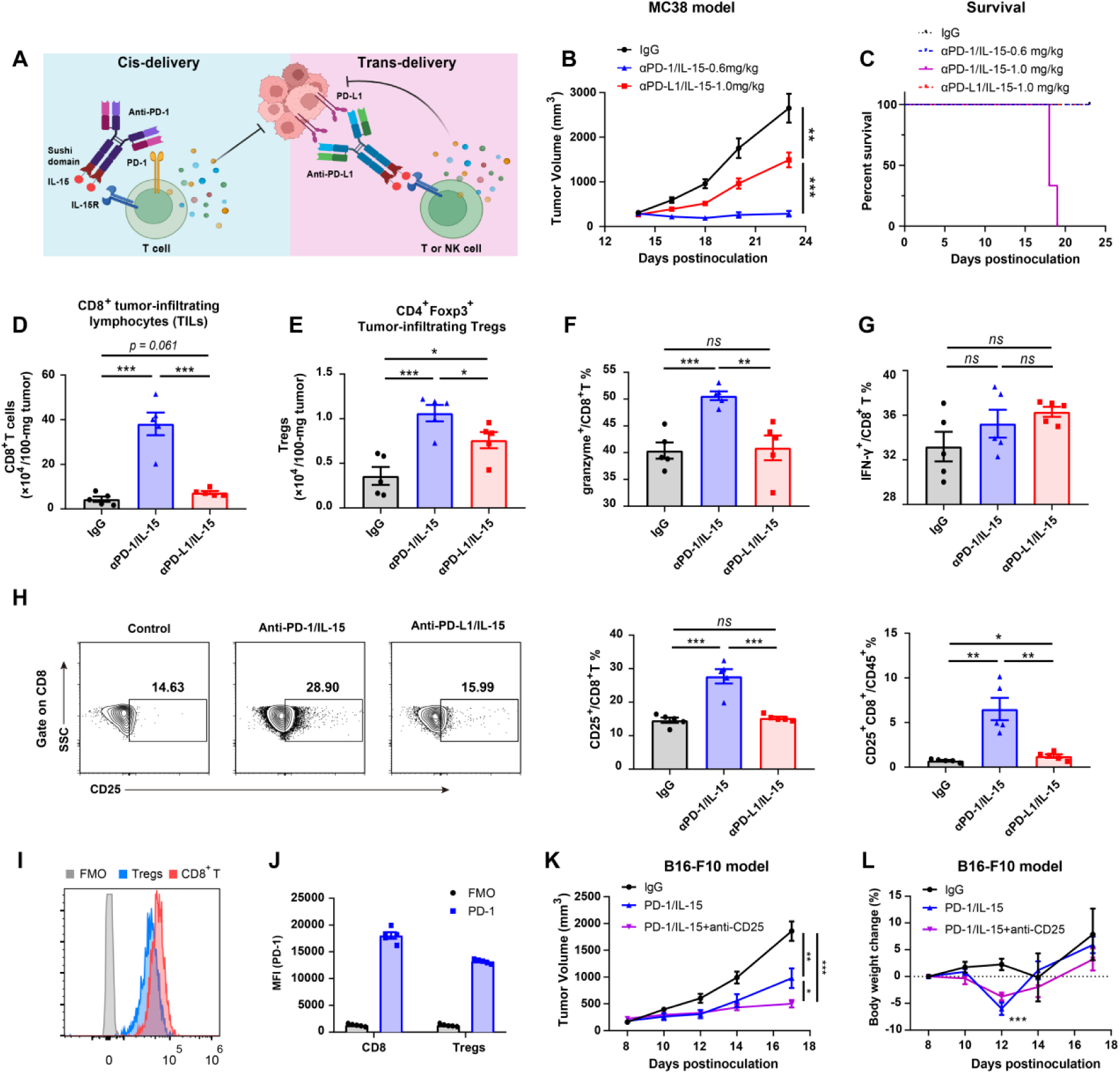
PD-1 cis-delivery shows superior antitumor effects compared to PD-L1 trans-presentation, but with higher toxicity. (A) Schematic diagram of antitumor mechanisms of cis-delivered and trans-delivered cytokines. (B and C) Female C57BL/6 mice were subcutaneously inoculated with 5 ×10^5^ MC38 tumor cells. The mice were then randomized into four groups based on tumor size, with treatment initiating when tumors reached 200 mm^3^. On days 14, 17, and 20, mice (n = 6) were intravenously injected with either IgG (1 mg/kg), αPD-L1/IL-15 (1 mg/kg), or αPD-1/IL-15 (0.6 mg/kg or 1 mg/kg). Tumor growth curves (B) and survival data (C) are shown. (D and E) Flow cytometry analysis of dissociated tumors (n = 5) was performed. The numbers of cells per 100 mg of tumor tissue were calculated for CD8^+^ tumor-infiltrating lymphocytes (TILs) (D) and tumor-associated Tregs (E). (F and G) Frequencies of granzyme B^+^CD8^+^ T cells (F) and IFN-γ^+^CD8^+^ T cells among CD8^+^ T cells (G) in the tumors are shown. (H) The percentages of CD25^+^CD8^+^ T cells among CD8^+^ T cells and CD45^+^ lymphocytes within tumors are shown, respectively. (I and J) Female C57BL/6 mice (n =5) were subcutaneously implanted with 5 × 10^5^ MC38 tumor cells. On days 18, mice were euthanized, and then tumor tissues were collected for flow cytometry analysis to evaluate PD-1 expression levels on tumor-infiltrating CD8^+^ T cells and Tregs. FMO, fluorescence minus one; MFI, mean fluorescence intensities. (K and L) Female C57BL/6 mice were subcutaneously inoculated with 4 × 10^5^ B16-F10 tumor cells. Tumor-bearing mice (n = 8) were intravenously treated with IgG or PD-1/IL-15 (0.6 mg/kg) on days 8, 11, and 14. For Tregs depletion, mice were intraperitoneally injected with 100 μg anti-CD25 on days 6, 9, and 12. Tumor growth curves (K) and body weight change of the tumor-bearing mice (L) are shown. All data represent mean ± SEM and are analyzed by one-way ANOVA or two-tailed Student’s t-test. \**p* < 0.05; \*\**p* < 0.01; \*\*\**p* < 0.001; ns, not significant (*p* > 0.05)

Flow cytometry analysis revealed that treatment with αPD-1/IL-15 (0.6 mg/kg) induced a significant increase in CD8^+^ tumor-infiltrating lymphocytes (TILs) compared to αPD-L1/IL-15 (1.0 mg/kg, Figure 1D and S1D). Unexpectedly, both αPD-1/IL-15 and αPD-L1/IL-15 treatments markedly increased the number of tumor-associated Tregs than IgG treatment, but αPD-1-mediated cis-delivery led to notably higher expansion rates than αPD-L1-mediated trans-delivery. These results demonstrate that targeted enrichment of IL-15 can stimulate Tregs, despite the conventional understanding that IL-15 has little effect on these cells (Figure 1E). Moreover, αPD-1/IL-15 treatment stimulated intratumoral CD8^+^ T cells to produce more granzyme B, indicating enhanced cytotoxicity. In contrast, αPD-L1/IL-15 treatment did not elevate the expression of either granzyme B or interferon γ (IFN-γ) in CD8^+^ T cells (Figure 1F and 1G). Additionally, in comparison with αPD-L1/IL-15, αPD-1/IL-15 treatment significantly expanded the CD25^+^CD8^+^ T cell subset which represents tumor-specific CD8^+^ T cells (TSTs), potentially accounting for its stronger antitumor activity (Figure 1H).

To evaluate the potential negative effects of αPD-1-mediated cytokine binding to Tregs, we detected PD-1 expression levels on CD8^+^ T cells and Tregs in the TME by flow cytometry. Both cell types exhibited high levels of PD-1 expression, although Tregs displayed slightly lower levels compared to CD8^+^ T cells (Figure 1I and 1J). To investigate the impact of Tregs on antitumor responses, we treated B16-F10 tumor-bearing mice with anti-CD25 depletion antibody. The results showed that the anti-CD25 antibody significantly enhanced the antitumor effect of αPD-1/IL-15, suggesting that Tregs diminish the antitumor activity of cis-delivering cytokines (Figure 1K). More importantly, since both TSTs and Tregs could be cleared by the anti-CD25 antibody, the depletion experiment revealed that αPD-1/IL-15 stimulated Tregs more than TSTs. It’s essential to block Tregs-induced immunosuppressive effects to further enhance antitumor efficacy. However, it is important to note that a dose of 0.6 mg/kg of αPD-1/IL-15 induced significant body weight loss, highlighting safety concerns associated with this molecule (Figure 1L). Collectively, compared to trans-presentation, cis-delivery of cytokine significantly enhances the proliferation and effector functions of CD8^+^ T cells, leading to a stronger antitumor effect. Nevertheless, it also exhibits greater adverse effects and notably stimulates Tregs.

### Engineering and characterization of a PD-1-targeted and conditionally-released IL-15 superagonist

To address the challenges associated with cis-delivery of cytokines, we need to achieve two primary objectives: first, to mitigate the severe immune-related adverse events (irAEs) caused by cytokines; and second, to minimize the stimulation of Tregs while preserving a robust stimulatory effect on CD8^+^ T cells. Considering the high biological activity of IL-15, we attempted to construct a heterodimer PSI-R, with one Fc chain fused to IL-15 and the other chain fused to the natural IL-15 receptor beta subunit to shield its activity. However, compared to the above described αPD-1/IL-15 (named PSI), PSI-R treatment only slightly reduced Mo7e cell proliferation, indicating that this strategy was not effective in masking IL-15 activity (Figure 2A, S2A and S2B). Drawing on strategies from structure-based drug design, we creatively employed steric hindrance from the antibody and sushi domain to conceal IL-15 activity. Thus, we engineered PIS containing an un-cleavable GS linker between anti-PD-1 and the IL-15/Sushi domain complex (ILR), resulting in significantly weaker proliferative activity than PSI in Mo7e cells (Figure S2C). Coincidentally, ILR has been reported as an IL-15 superagonist that is more potent and stable than wild-type IL-15(21). Therefore, we replaced the uncleavable linker between anti-PD-1 and ILR with a matrix metalloproteinase (MMP) cleavable linker to construct a conditionally-cleavable anti-PD-1/IL-15 (named PMIS). When anti-PD-1 directs ILR to the TME, PMIS would be cleaved by tumor-associated proteases, releasing ILR to freely stimulate all target cells, including CD8^+^ T cells and NK cells, instead of being bound to Tregs by anti-PD-1 (Figure 2A). This approach is expected to minimize the negative impact of Tregs on the antitumor immune response.

**Figure 2.**
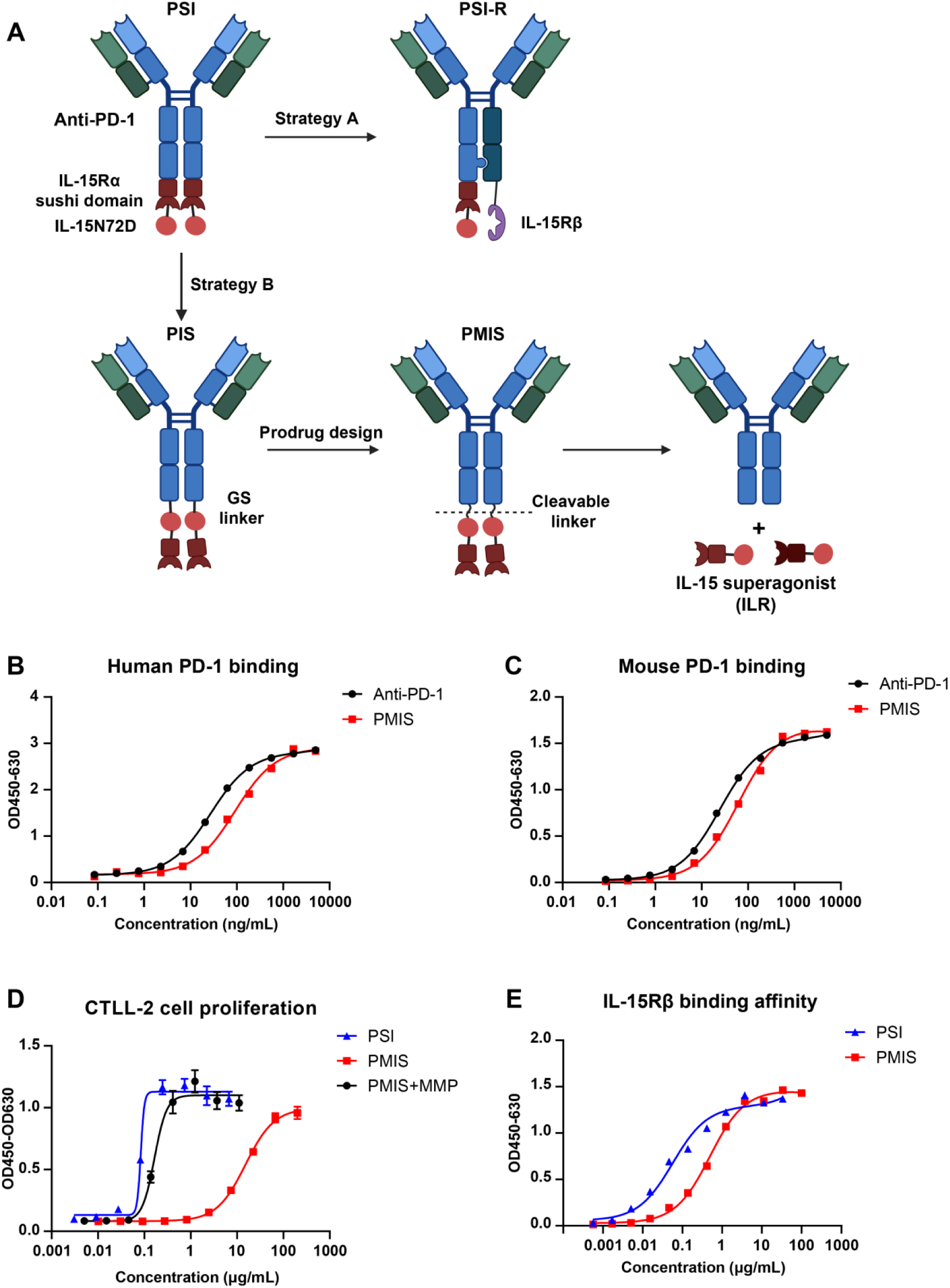
Design and *in vitro* characterization of conditionally activated anti-PD-1/IL-15. (A) Design diagram of conditionally-activated and cis-delivery of IL-15 superagonist. (B and C) Binding of anti-PD-1 and PMIS to plate-bound human or mouse PD-1 (n = 2 technical replicates). Data were analyzed using the one site-total to calculate the EC_50_ values. (D) The proliferative potential of PMIS and MMP-cleaved PMIS was compared with PSI in mouse CTLL-2 cells (n = 3 technical replicates). Data were analyzed using the four-parameter-fit logistic equation to calculate the EC_50_ values. Data was shown as mean ±SD. (E) Binding of PSI and PMIS to plate-bound human IL-15Rβ (n = 2 technical replicates). Data were analyzed using the one site-total to calculate the affinity values.

ELISA results demonstrated that PMIS bound to human PD-1 with a profile similar to that of anti-PD-1 (EC_50_ = 91.34 and 27.07 ng/mL or 0.44 nM and 0.17 nM, respectively) (Figure 2B), and it also exhibit comparable affinity for mouse PD-1 (EC_50_ = 61.72 and 24.75 ng/mL or 0.30 and 0.16 nM, respectively) (Figure 2C). This indicates the binding affinity of the anti-PD-1 portion of PMIS was preserved.

SDS-PAGE analysis revealed that PMIS, but not PIS, can be cleaved after incubation with MMP (Figure S2D). PMIS demonstrated a 177-fold reduction in proliferative activity than PSI in CTLL-2 cells (EC_50_ = 15.19 μg/mL or 73.81 nM versus 0.084 μg/mL or 0.42 nM). Upon cleavage, it restored its activity by more than 90-fold (EC_50_ = 0.16 μg/mL or 0.80 nM) (Figure 2D). PMIS also significantly reduced proliferative activity compared to PSI in Mo7e cells, and following cleavage, it notably restored proliferative activity, similar to the effect observed in CTLL-2 cells (Figure S2E). PMIS also showed a decreased IL-15Rβ binding affinity compared with PSI (490.6 ng/mL or 2.38 nM versus 56.51 ng/mL or 0.28 nM) due to IL-15 masking, which accounts for its weaker proliferative activity (Figure 2E). In short, PMIS demonstrated significantly diminished stimulatory activity, but once cleaved, the released ILR can effectively restore its immunostimulatory activity.

### PMIS minimizes systemic irAEs compared to PSI

To determine whether our strategy can reduce the irAEs caused by IL-15, healthy Balb/c mice were administered either 1 mg/kg of PSI, or 5 or 10 mg/kg of PMIS (representing 5-fold and 10-fold molar doses of PSI at 1 mg/kg, respectively). It was observed that mice treated with PSI experienced significant body weight loss compared to the PBS treatment, while mice treated with 5 or 10 mg/kg of PMIS maintained their body weight (Figure 3A). After two treatments, 40% of mice receiving PSI died within 6 days, whereas none of mice administered PMIS lost weight or died (Figure 3B). PSI at a dose of 1 mg/kg increased plasma IFN-γ levels in a large extent at 8 and even 48 hours post-treatment, while no significant elevation in IFN-γ levels was observed following 5 mg/kg of PMIS treatment. Although plasma IFN-γ levels showed a slight increase 8 hours after treatment with 10 mg/kg of PMIS, they returned to baseline levels by 48 hours (Figure 3C). Unlike PSI, both 5 and 10 mg/kg of PMIS did not elevate alanine aminotransferase (ALT) and aspartate aminotransferase (AST) levels in blood (hepatotoxicity markers) (Figure 3D). These results demonstrate a significant reduction in irAEs with PMIS compared to PSI.

**Figure 3.**
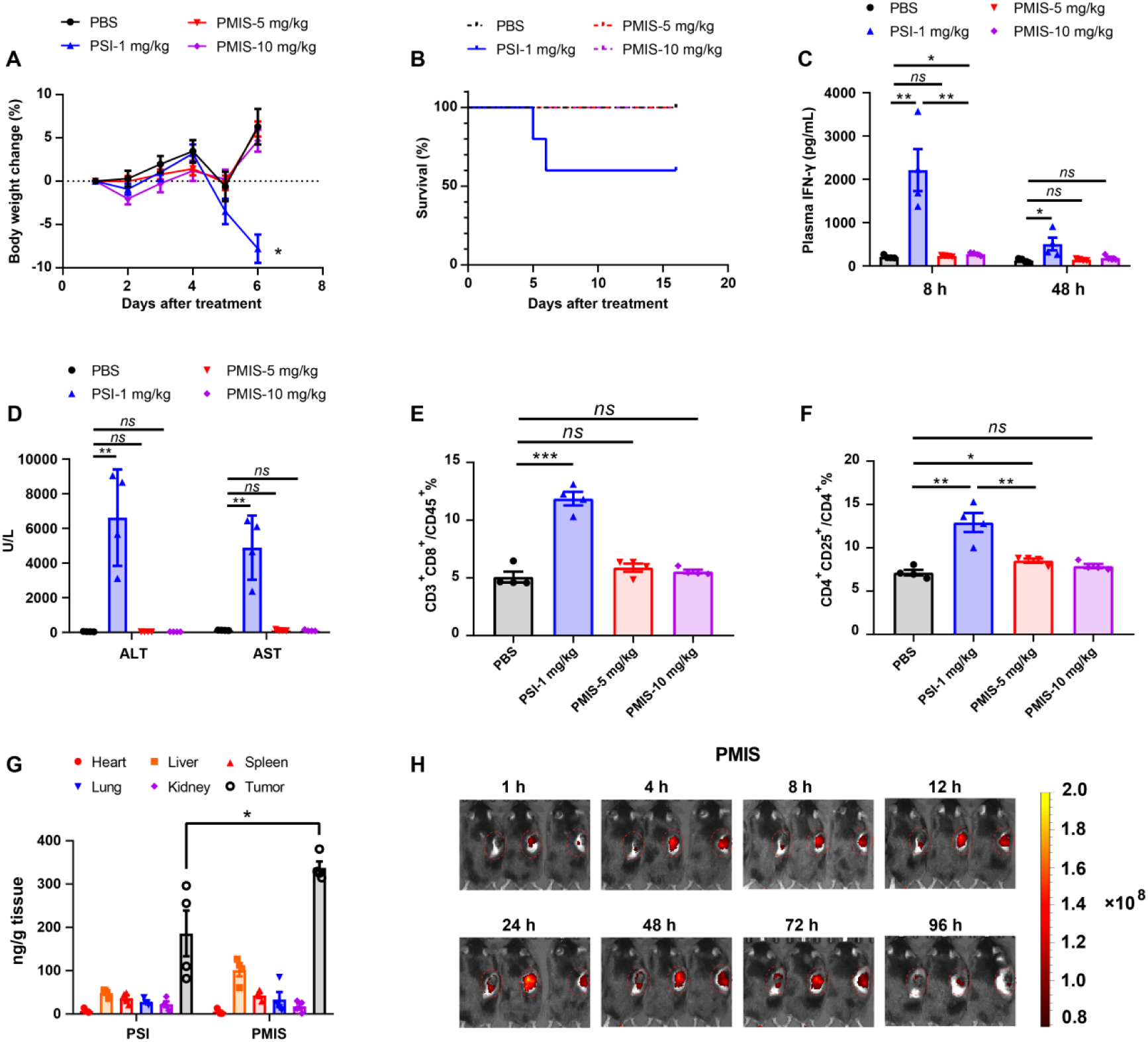
PMIS reduces systemic toxicity associated with IL-15. (A and B) Female Balb/c mice were intravenously injected with PBS, PSI (1.0 mg/kg), or PMIS (5 or 10 mg/kg) on days 1 and 4, with body weight and survival monitored (n = 5). (C) Female Balb/c mice were intravenously injected with PBS, PSI (1.0 mg/kg), or PMIS (5 or 10 mg/kg) on days 1 and 4 (n = 4). Blood were collected at 8 and 48 hours after the second treatment, and plasma IFN-γ levels were measured using ELISAs. (D) Blood were collected at 48 hours after the second treatment (n = 4). The levels of ALT and AST in the plasma were quantified. (E and F) Spleens of mice were extracted after euthanasia on day 6. The percentages of splenic CD8^+^ T cells and Tregs are shown for populations of CD45^+^ lymphocytes, respectively (n = 4). (G) B16-F10 tumor-bearing mice (n = 4) were intravenously injected with 3 mg/kg PSI or PMIS, and tissues were collected at 48 h post injection, respectively. The concentrations of PSI or PMIS were measured using ELISA. (H) Cy5.5-labeled PMIS (1 mg/kg) was intravenously injected into RM-1 tumor-bearing mice. PMIS accumulation in the tumors was tracked by the IVIS Spectrum *in vivo* imaging system (n = 3). All data represent mean ± SEM and are analyzed by one-way ANOVA or two-tailed Student’s t-test. \**p* < 0.05; \*\**p* < 0.01; \*\*\**p* < 0.001; ns, not significant (*p* > 0.05)

We subsequently employed flow cytometry to examine the effects of PMIS on peripheral T cells. Intriguingly, PMIS treatment neither increased the percentage of CD8^+^ T cells nor elevated the proportion of CD4^+^CD25^+^ T cells, indicating that PMIS, in its un-cleaved form, lacks stimulatory activity on T cells, including both CD8^+^ T cells and Tregs (Figure 3E, 3F, and S3A). Furthermore, we investigated the tissue distribution of PSI and PMIS. The results revealed that while both PSI and PMIS effectively targeted tumors, PMIS showed an even greater targeting ability (Figure 3G). To trace the fate of PMIS at the tumor site, we conjugated it with a Cy5.5-maleimide tracer and injected it into tumor-bearing mice for whole-body imaging at various time points. The bioluminescence data indicated a gradual increase in tumor fluorescence intensity, peaking at approximately 12 h, keeping stable between 12 and 72 h, and then beginning to decline after 72 h (Figure 3H).

### PMIS enhances antitumor efficacy while maintaining a favorable safety profile

To explore whether PMIS exhibits antitumor activity in a dose-dependent manner, we first evaluated its effects at various doses using the B16-F10 cold tumor model, which is characterized by high MMP expression(22). Our findings revealed that PMIS demonstrated significant antitumor efficacy across all tested doses, with a notably stronger antitumor effect at a dosage of 5 mg/kg compared to 2 mg/kg. No statistically significant difference was observed between the 5 mg/kg and 10 mg/kg groups (Figure 4A and S4A). In addition, PMIS treatment at various doses did not result in body weight loss or spleen weight increase of mice, underscoring its favorable safety profile (Figure S4B and S4C). Moreover, histopathological section analysis illustrated that PMIS treatment (10 mg/kg) did not induce liver or kidney damage (Figure S4D). Furthermore, we compared the antitumor effects of PMIS with PSI and PIS in the B16-F10 tumor model. PIS and PSI demonstrated comparable antitumor effects, but were significantly weaker than that of PMIS (Figure 4B). PMIS notably extended the lifespan of the mice, with 10% achieving long-term survival (Figure 4C and S4E). In addition, PSI induced obvious body weight loss, suggesting that its antitumor effects come at the expense of severe toxicity (Figure S4F). Moreover, PSI treatment induced a significant increase in spleen weight compared to PMIS treatment, indicating that PMIS has improved tumor-targeting capabilities and reduced peripheral immunostimulatory activity (Figure S4G).

**Figure 4.**
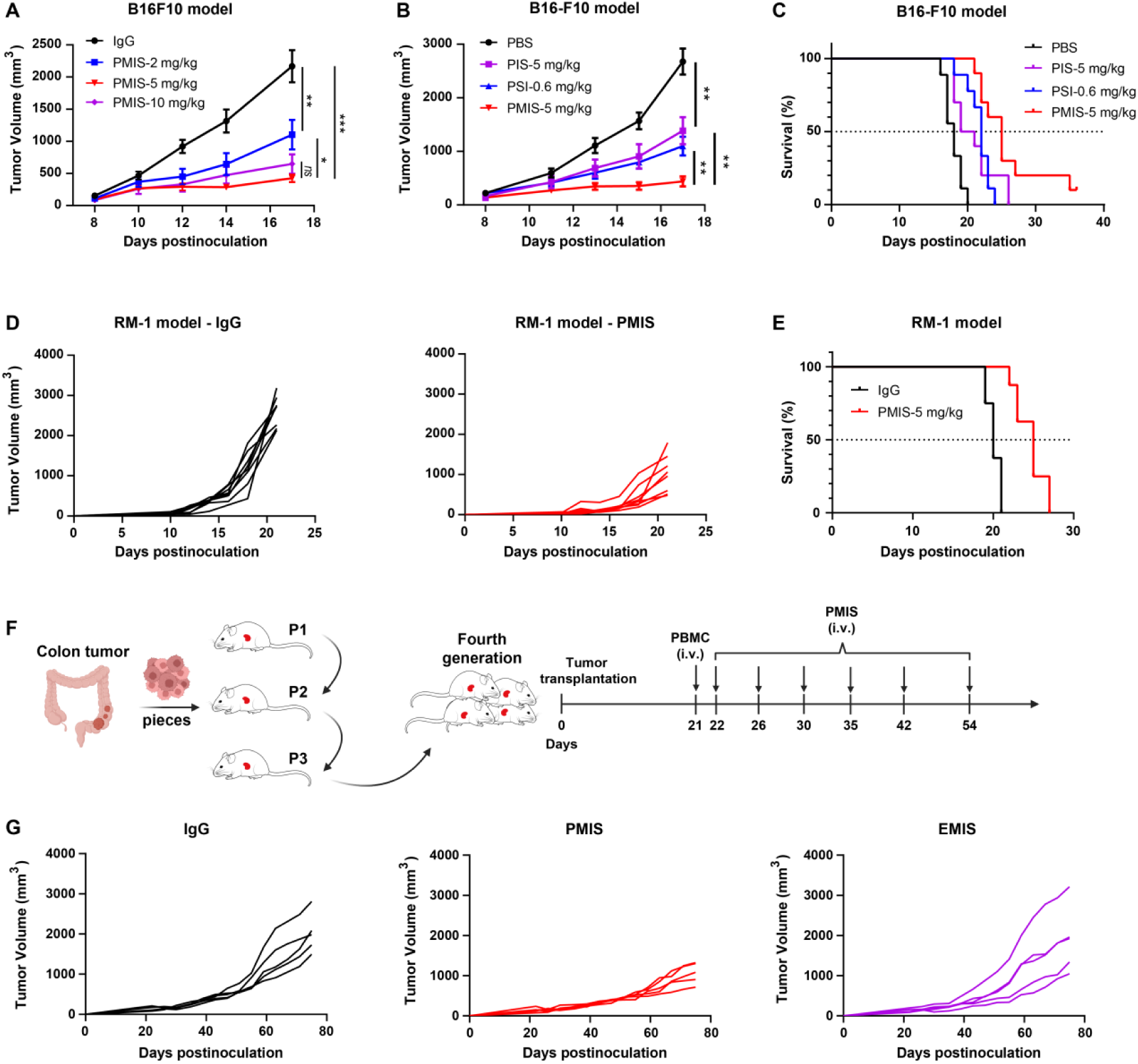
PMIS demonstrates superior antitumor activity in cold and advanced tumors. (A) B16-F10 tumor cells (4×10^5^) were subcutaneously implanted into female C57BL/6 mice. Mice were then randomized into four groups, and treatment initiated when tumors reached 50-100 mm^3^. On days 8, 11, and 14 (n = 7), mice were intravenously injected with IgG control (5 mg/kg) or PMIS (2, 5, or 10 mg/kg). Tumor volumes were measured as indicated. (B and C) B16-F10 tumor cells (4×10^5^) were subcutaneously implanted into female C57BL/6 mice. Mice were then randomized into four groups, and treatment initiated when tumors reached 50-100 mm^3^. On days 8, 11, and 14 (n = 9-10), mice were intravenously injected with PBS control, PIS (5 mg/kg), PSI (0.6 mg/kg), or PMIS (5 mg/kg). Tumor growth curves (B) and survivals (C) are plotted. (D and E) RM-1 tumor cells (5×10^5^) were subcutaneously implanted into male C57BL/6 mice. Mice were then randomized into two groups, and treatment initiated when tumors reached 50-100 mm^3^. On days 10, 13, and 16 (n = 8), mice were intravenously injected with IgG control or PMIS (5 mg/kg). Tumor growth curves (D) and survival (E) are shown. (F and G) Schematic diagram of establishment process of the PDX model of colorectal carcinoma (F). Tumor-bearing mice were randomized into three groups (n = 5), and treatment initiated when tumors reached 100-150 mm^3^. Mice received intravenously 4 × 10^6^ fresh human peripheral blood mononuclear cells (PBMCs) on day 21, followed by intravenous administration of IgG (5 mg/kg), PMIS (5 mg/kg), or EMIS (5 mg/kg) on days 22, 26, 30, 35, 44, and 54. Tumor growth curves are shown (G). All data represent mean ± SEM and are analyzed by one-way ANOVA or two-tailed Student’s t-test. \**p* < 0.05; \*\**p* < 0.01; \*\*\**p* < 0.001; ns, not significant (*p* > 0.05)

Next, we utilized the murine RM-1 prostate cold tumor model to further evaluate the antitumor potential of PMIS. The results demonstrated that PMIS significantly inhibited tumor growth and effectively prolonged the survival of RM-1 tumor-bearing mice (Figure 4D, 4E, and S4H). Additionally, PMIS did not induce weight loss in these mice (Figure S4I).

Patient-derived xenograft (PDX) model excels at reflecting the characteristics of cancer and simulating tumor progression and evolution in human patients, thereby yielding the most convincing preclinical results. We established a PDX model of colorectal carcinoma to investigate the antitumor potential of PMIS (Figure 4F). To evaluate the critical role of anti-PD-1 mediated cis-delivery in antitumor effects, we constructed a control prodrug targeting EGFR, named EMIS (Figure S4J). According to the results, PMIS demonstrated remarkable antitumor efficacy in the PDX model, underscoring its considerable potential for clinical translation. However, EMIS treatment showed no significant antitumor effects, highlighting the essential role of anti-PD-1 and its cis-delivery of IL-15 in tumor control (Figure 4G). Furthermore, PMIS treatment significantly reduced Ki67 expression in tumors compared to the other two treatments, indicating a decreased proliferative capacity of tumors (Figure S4K). In summary, tumor-conditional cis-delivery of IL-15 superagonist demonstrated enhanced antitumor activity while significantly minimizing systemic toxicity.

### PMIS significantly increases CD25^+^CD8/Treg ratio within tumors

To further explore the mechanisms underlying the superior antitumor effects of PMIS, we performed flow cytometry analysis of the dissociated B16-F10 tumors. Compared to the PBS control, treatment with PIS, PSI, and PMIS all increased the proportions and numbers of intratumoral CD8^+^ T cells (Figure 5A, S5A and S5B). Notably, treatment with PMIS resulted in a higher percentage of CD8^+^ TILs compared to PSI. Furthermore, PMIS treatment significantly increased the proportion of CD25^+^CD8^+^ TILs relative to PSI treatment (Figure 5B). Besides, compared with PBS control, all treatments significantly raised the numbers of Tregs, while only causing a slight increase in the proportions of Tregs (Figure S5C and S5D). Moreover, PMIS treatment resulted in markedly higher CD25^+^CD8/Treg and CD8/Treg ratios within the tumor compared to PSI treatment, which could clarify why PMIS exhibited much stronger antitumor efficacy than PSI (Figure 5C and S5E).

**Figure 5.**
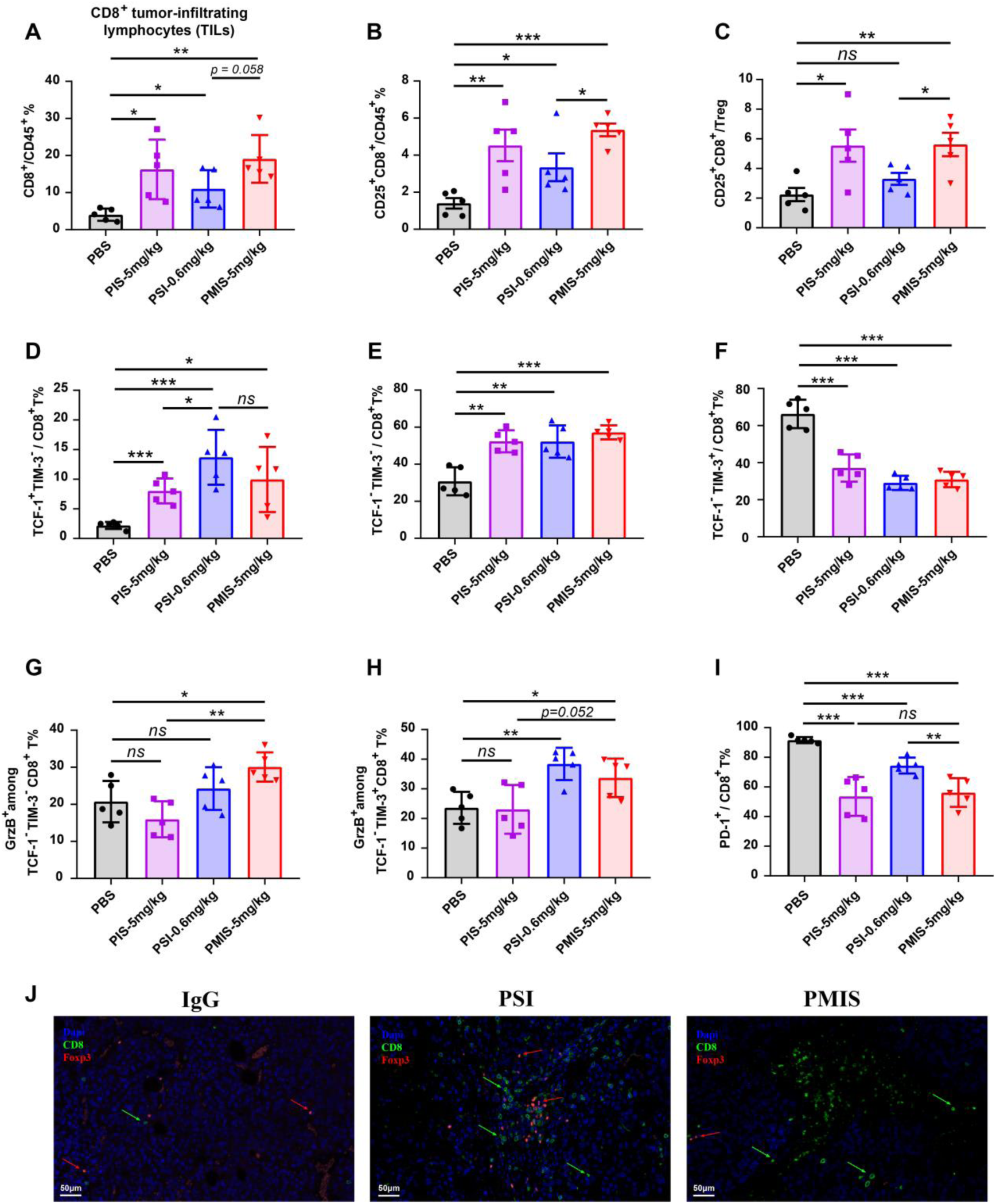
PMIS drives exhausted CD8^+^ T cell effector function to enhance antitumor activity. B16-F10 tumor cells (4×10^5^) were subcutaneously implanted into female C57BL/6 mice. Mice were then randomized into four groups, and treatment initiated when tumors reached 50-100 mm^3^. On days 8, 11, and 14, mice (n = 5) were intravenously injected with PBS control, PIS (5 mg/kg), PSI (0.6 mg/kg), or PMIS (5 mg/kg). On day 17, mice were euthanized, and tumors were removed for flow cytometry analysis and immunofluorescence staining. (A and B) The percentages of intratumoral CD8^+^ T cells and CD25^+^CD8^+^ T cells within the CD45^+^ lymphocyte populations are shown, respectively (n = 5). (C) CD25^+^CD8/Treg ratios are calculated. (D-F) Percentages of TCF-1^+^TIM-3^-^ (D), TCF-1^-^TIM-3^-^ (E), and TCF-1^-^TIM-3^+^ (F) subpopulations among CD8^+^ TILs are shown. (G and H) Frequencies of granzyme B^+^CD8^+^ T cells among TCF-1^-^TIM-3^-^ (G) and TCF-1^-^TIM-3^+^ (H) are shown, respectively. (I) Frequency of PD-1^+^CD8^+^ T cells among CD8^+^ T cells are shown. (J) Immunofluorescence staining for CD8 (green) and Foxp3 (red) of tumor tissues (scale bar, 50 μm; n = 2). All data represent mean ± SEM and are analyzed by one-way ANOVA or two-tailed Student’s t-test. \**p* < 0.05; \*\**p* < 0.01; \*\*\**p* < 0.001; ns, not significant (*p* > 0.05)

Although PIS and PMIS induced comparable CD8/Treg ratios, the antitumor effect of PIS was significantly weaker than that of PMIS. To further understand this discrepancy, we analyzed the changes in the various subsets of CD8^+^ T cells and their effector functions. The results indicated that all treatments markedly elevated the percentages of TCF-1^+^TIM-3^-^ (stem-like) and TCF-1^-^TIM-3^-^ (effector) populations, while decreasing the frequency of TCF-1^-^TIM-3^+^ (terminally exhausted) cells within the CD8^+^ TILs (Figure 5D-5F). Additionally, both PMIS and PSI treatments increased the production of granzyme B, a key cytotoxic molecule, in terminally exhausted CD8^+^ T cells, whereas only PMIS upregulated granzyme B expression in effector CD8^+^ T cells (Figure 5G and 5H). Of note, PIS treatment did not enhance granzyme B expression in either effector or terminally exhausted CD8^+^ T cells, which may partially explain why the antitumor activity of PIS is inferior to that of PMIS (Figure 5G and 5H). Moreover, compared to PSI, PMIS treatment significantly downregulated PD-1 expression in CD8^+^ TILs, suggesting that PMIS treatment can more effectively reverse intratumoral CD8^+^ T cell exhaustion (Figure 5I). Immunofluorescence analysis further revealed that PMIS treatment significantly increased the number of intratumoral CD8^+^ T cells and the CD8/Treg ratio (Figure 5J). Altogether, PMIS treatment can enhance the intratumoral CD25^+^CD8/Treg ratio, reverse T cell exhaustion, and improve the effector function of CD8^+^ T cells, resulting in enhanced antitumor effects.

### The antitumor activity of PMIS depends on pre-existing intratumoral CD25^+^CD8^+^ T cells

To further elucidate the differences in antitumor efficacy between PSI and PMIS, we conducted RNA sequencing (RNA-seq) on tumor tissues. Compared to PSI, PMIS induced a greater infiltration of immune cells (Figure 6A). Gene set enrichment analysis revealed significant enrichment of pathways and gene expression associated with leukocyte-mediated cytotoxicity (Figure S6A). Consistent with the flow cytometry data shown in Figure 5, PMIS significantly increased intratumoral CD8^+^ T cells and slightly elevated Foxp3 expression compared to PSI (Figure 6B). Moreover, PMIS treatment notably enhanced T cell-mediated cytotoxicity relative to PSI (Figure 6C and S6B). Overall, PMIS not only expanded intratumoral CD8^+^ T cells but also boosted their cytotoxicity, leading to a stronger antitumor immune response.

**Figure 6.**
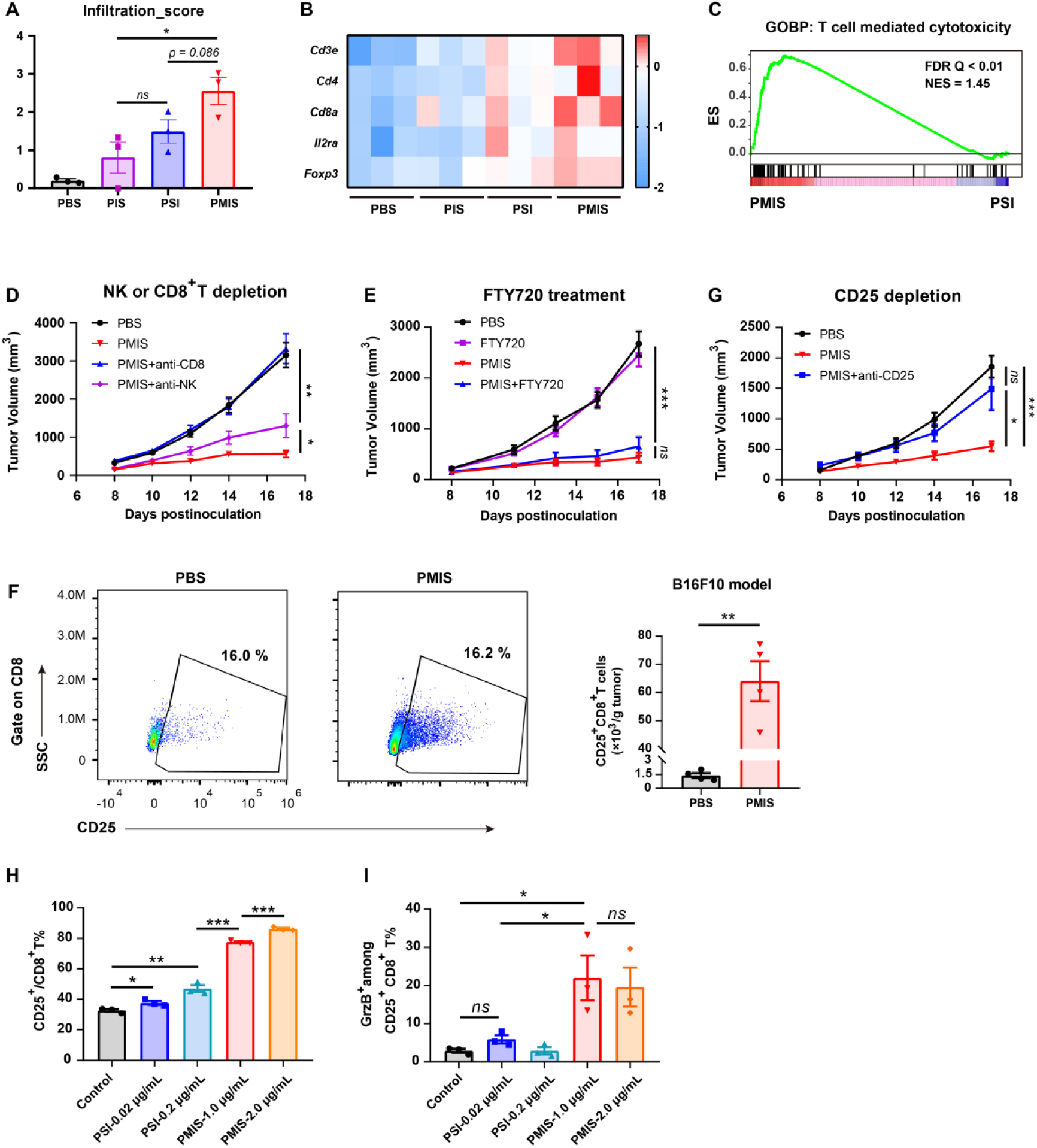
Intratumoral CD25^+^CD8^+^T cells mediate the antitumor efficacy of PMIS. (A-C) Tumor tissues of B16-F10-bearing mice treated as described in Figure 5 were extracted for RNA-seq analysis (n = 3). Immune infiltration analysis was performed using immuCellAI (A). The heatmap depicts the alterations in gene expression of T cell markers in response to different treatments (B). Gene set enrichment analysis was performed for T cell mediated cytotoxicity after PMIS or PSI treatment (C). ES, enrichment score; FDR, false discovery rate; NES, normalized enrichment score. (D) B16-F10 tumor-bearing mice (n = 6) were intravenously treated with PBS or PMIS (5 mg/kg) on days 8, 11, and 14. Mice were intraperitoneally injected with 200 μg of anti-NK1.1 antibody or 200 μg of anti-CD8α antibody on days 6, 8, and 12. Tumor growth curves were plotted. (E) B16-F10 tumor-bearing mice (n = 7) were intravenously treated with PBS or PMIS (5 mg/kg) on days 8, 11, and 14. Then FTY720 (1.25 mg/kg) was administered intraperitoneally on days 7, 9, 11, 13, and 15 to block T cells egress from the lymph nodes. Tumor growth curves were shown. (F) On day 17, mice (n = 4) were euthanized, and tumors were removed for flow cytometry analysis. Numbers of CD25^+^CD8^+^ T cells per gram of tumor tissue were calculated. (G) Female C57BL/6 mice were subcutaneously inoculated with 4×10^5^ B16-F10 cells. Mice were then randomized into three groups, and treatment initiated when tumors reached 50-100 mm^3^. On days 8, 11, and 14, mice (n = 8) were intravenously injected with PBS control or PMIS (5 mg/kg). For CD25^+^ T cell depletion, mice were intraperitoneally injected with 100 μg of anti-CD25 on days 6, 9, and 12. Tumor growth curves are shown. (H and I) Mouse splenic T cells were isolated, activated, and then incubated with PBS, PSI (0.02 or 0.2 μg/mL), or PMIS (1.0 or 2.0 μg/mL). After 48 h, T cells were collected for flow cytometry analysis (n = 3). All data represent mean ± SEM and are analyzed by one-way ANOVA or two-tailed Student’s t-test. \**p* < 0.05; \*\**p* < 0.01; \*\*\**p* < 0.001; ns, not significant (*p* > 0.05)

Considering that IL-15 can activate both T and NK cells, we used specific depletion antibodies to remove CD8^+^ T or NK cells in B16-F10 tumor-bearing mice to determine their respective contributions to PMIS’s antitumor effects. The therapeutic effect of PMIS was abolished in mice with CD8^+^ T cells depletion and less affected by NK-cell depletion (Figure 6D). We then used FTY720 to further investigate whether pre-existing immune cells within the tumor or recruited cells contribute to PMIS’s antitumor efficacy(23). Blocking the egress of T cells from lymphoid tissues into the tumor did not diminish the antitumor effects of PMIS, indicating that pre-existing intratumoral T cells are sufficient for PMIS therapy (Figure 6E). Furthermore, PMIS treatment markedly increased the number of intratumoral CD25^+^CD8^+^ T cells (Figure 6F). To assess their role, we conducted a depletion experiment using anti-CD25 antibody. Intriguingly, anti-CD25 treatment completely compromised the antitumor effects of PMIS, indicating that CD25^+^CD8^+^ T cells are crucial mediators of its antitumor activity (Figure 6G). To further validate the stimulatory activity on CD25^+^CD8^+^ T cells, we incubated the fusions with mouse primary splenic T cells. We find that the cleaved PMIS significantly stimulated the proliferation of CD25^+^CD8^+^ T cells and enhanced their effector function compared with PSI, which explains the superior antitumor efficacy of PMIS (Figure 6 H and 6I). In summary, the therapeutic effect of PMIS was primarily driven by intratumoral CD25^+^CD8^+^ T cells, whereas NK cells and the influx of newly recruited T cells into the TME also play important roles, they are less critical to the antitumor immunity induced by PMIS. The above findings also indicate that CD25^+^CD8^+^ T cells play an important role in the antitumor effect of IL-15-based immunotherapy.

### Cis-delivery of IL-15 potently suppresses orthotopic cold carcinoma and its metastasis, enhanced by combination with chemotherapy

Triple-negative breast cancer (TNBC), which accounts for approximately 15-20% of breast malignancies, is characterized by early recurrence and repetitious metastasis(24). For a long time, chemotherapy was the only systemic treatment available for TNBC, but the responses were often short-lived(25). In recent years, immunotherapies have been explored for TNBC, but they demonstrate only modest efficacy, with a small proportion of patients benefiting from this approach(26). Therefore, improved therapies are urgently needed to extend survival in TNBC patients, especially those with metastasis.

We utilized the 4T1 orthotopic breast cancer spontaneous metastasis mouse model to evaluate the antitumor effects of PMIS, both as a standalone treatment and in combination with doxorubicin, a widely used chemotherapy agent for TNBC. The results revealed that both PMIS and doxorubicin treatments significantly inhibited *in situ* tumor growth, and the combination therapy exhibited synergistic antitumor efficacy (Figure 7A). Compared to IgG treatment, PMIS notably controlled tumor lung metastasis, whereas doxorubicin did not show a similar effect. The combination treatment demonstrated enhanced suppression of tumor metastasis compared to PMIS monotherapy, though this difference was not statistically significant (Figure 7B).

**Figure 7.**
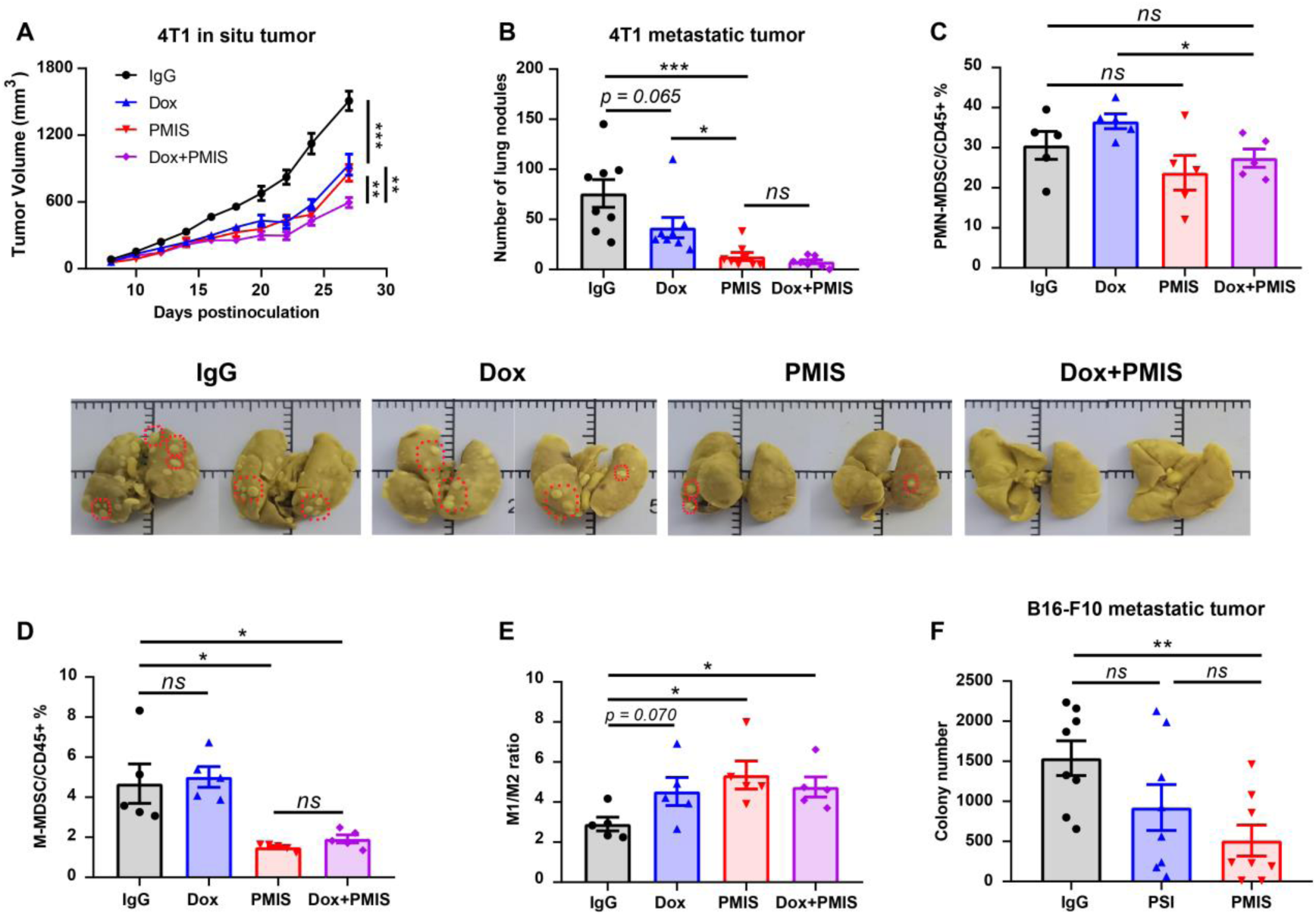
PMIS effectively inhibits both *in situ* cancer and its metastasis, enhanced by combination with chemotherapy. (A and B) Female Balb/c mice were orthotopically inoculated with 5×10^5^ 4T1 cells in the fourth mammary fat pad. Mice were then randomized into four groups, and treatment initiated when tumors reached 50-100 mm^3^. Tumor-bearing mice (n = 8) were intravenously treated with IgG or PMIS (5 mg/kg) on days 8 and 13. Mice were intravenously treated with doxorubicin (5 mg/kg) on days 9 and 15. Tumor growth curve was shown (A). On day 27, after euthanizing the mice, remove their lungs, rinse with PBS, and then immerse in Bouin solution in the dark for 12 hours. The colonies on the lungs were counted (B). (C and D) The percentages of intratumoral PMN-MDSCs and M-MDSCs within CD45^+^ lymphocytes are shown, respectively (n = 5). (E) The ratios of M1/M2 were calculated (n = 5). (F) B16-F10 tumor cells (4×10^5^) were intravenously implanted into the female C57BL/6 mice. On days 6, 9, and 12, mice (n = 8) were intravenously injected with IgG control (5 mg/kg), PSI (0.6 mg/kg), or PMIS (5 mg/kg). The mice were sacrificed on day 15, and lung colonies were counted. All data represent mean ± SEM and are analyzed by one-way ANOVA or two-tailed Student’s t-test. \**p* < 0.05; \*\**p* < 0.01; \*\*\**p* < 0.001; ns, not significant (*p* > 0.05)

IL-15 is a pleiotropic cytokine capable of regulating various immune cells, including T and NK cells, to ameliorate antitumor responses(27). Given its role in modulating the TME, we specifically investigated its effects on myeloid-derived suppressor cells (MDSCs) and macrophages, which are crucial in tumor immunosuppression. The results showed that PMIS treatment slightly reduced the percentage of polymorphonuclear (PMN)-MDSC cells and significantly decreased monocytic (M)-MDSC cells compared to the control group. Doxorubicin treatment did not have a significant impact on either subset of MDSCs (Figure 7C and 7D). Moreover, PMIS significantly increased the M1/M2 ratio of tumor-associated macrophages compared to IgG, suggesting its ability to repolarize macrophages to enhance antitumor immunity (Figure 7E). Furthermore, in a murine B16-F10 metastatic melanoma model, PMIS treatment markedly reduced lung colony numbers compared to the IgG control, whereas PSI did not (Figure 7F). Collectively, the remarkable antitumor efficacy of PMIS presents a new option for the treatment of refractory and metastatic cold tumor.

## Discussion

In the early stages of immunocytokines development, attempts have been made to selectively deliver cytokines to the TME via fusing them to antibodies targeting tumor-associated antigens or tumor stroma(28). This approach aimed to concentrate cytokines at the tumor site to achieve two primary goals: reducing the toxic side effects of cytokine and enhancing synergistic antitumor efficacy. However, these strategies have been proven to be less efficient than anticipated(29). Immunocytokines based on trans-presentation have been in development for decades, yet none have received market approval(4). Utilizing anti-PD-1 to deliver cytokines to tumor-infiltrating PD-1^+^CD8^+^ T cells in a cis-presentation manner has attracted increasing attention in recent years, as it can effectively reactivate intratumoral exhausted CD8^+^ T cells, resulting in potent antitumor activity(4). In this study, for the first time, we performed a direct comparison of cis-delivery and trans-delivery of cytokine to evaluate their differences in antitumor effects and safety profiles. Our findings revealed that anti-PD-1/IL-15 exhibited significantly stronger antitumor effects than anti-PD-L1/IL-15, markedly expanding tumor-infiltrating CD8^+^ T cells and enhancing their effector functions, but it also exhibited increased toxicity and potent Tregs stimulatory activity.

Given the high PD-1 expression on intratumoral Tregs, there is always a risk that anti-PD-1-mediated cis-presentation of cytokine could stimulate these cells. As shown in Figure 1E, the effect of IL-15 on Tregs has traditionally been regarded as negligible, while PD-1 cis-delivery significantly enhances this effect compared to PD-L1 trans-delivery. CD25^+^ cells depletion experiments further demonstrated that the antitumor effect of αPD-1/IL-15 could be improved by eliminating immunosuppressive Tregs.

IL-2 and IL-15 are the most extensively studied cytokines for cis-delivery. To selectively activate IL-2Rβγ-expressing CD8^+^ T and NK cells over IL-2Rαβγ-expressing Tregs, most IL-2-based cancer therapeutics currently being evaluated utilize an IL-2Rβγ-biased agonism strategy(30–32). However, recent studies have demonstrated that engaging IL-2Rα is critical for the antitumor effect of systemic IL-2 therapy. Both Innovent and Regeneron’s research findings suggested that although IL-2Rβγ-biased IL-2 can systemically expand CD8^+^ T and NK cells, these expanded cells are not tumor specific(18, 33). This lack of specificity may explain why the antitumor effects of IL-2 Rβγ-biased agonists are less significant than wild-type IL-2. Moreover, Innovent’s findings revealed that IL-2Rα-biased agonist can invigorate tumor-infiltrating CD25^+^CD8^+^ T cells. But they did not discuss the expansion of Tregs by IL-2Rα-biased agonists. This suggests that the antitumor effects of the cis-delivery cytokines depend on the balance of CD25^+^CD8^+^ T cells and Tregs.

To maximize the stimulation of CD25^+^CD8^+^ T cells while minimizing the activation of Tregs, we introduced a cleavable linker between anti-PD-1 and ILR to construct PMIS. Upon reaching the tumor site, immunostimulatory ILR is released through cleavage by tumor-associated proteases, rather than being restricted to CD8^+^ T cells or Tregs mediated by anti-PD-1. This design allows ILR to act freely on various immune cells, thereby reducing its stimulatory effect on Tregs and mitigating their detrimental impact. Animal experiments demonstrated that PMIS significantly improved antitumor efficacy by achieving a markedly higher CD25^+^CD8/Treg ratio compared to other treatments. In contrast to PSI, targeted depletion of CD25^+^ T cells using anti-CD25 antibodies completely compromised the efficacy of PMIS, highlighting its dependence on the specific stimulation of CD25^+^CD8^+^ T cells. Additionally, we constructed PIS, which also has sterically masked ILR like PMIS but cannot be cleaved. PIS showed a similar safety profile and intratumoral CD25^+^CD8 /Treg ratio, but demonstrated weaker antitumor effects. Further experiment revealed that in comparison with PIS, PMIS stimulated exhausted CD8^+^ T cells more potently and enhanced their cytotoxicity, attributed by the release of ILR. Collectively, PMIS exhibited unprecedented safety and efficacy in comparison with all other strategies for cis-delivery IL-15-based immunocytokines.

Since anti-PD-1-mediated cis-delivery can redirect cytokine to intratumoral PD-1^+^CD8^+^ T cells, anti-PD-1-based immunocytokines can be effectively combined with other therapeutics such as chemotherapy, chimeric antigen receptor (CAR)-T cells, and so on. In our study, the combination of PMIS and doxorubicin exhibited significant synergistic antitumor efficacy in the TNBC 4T1 model (Fig. 7A&B). This synergy is likely due to doxorubicin inducing tumor cell death and releasing tumor antigens, which prime the immune system, while PMIS enhances the immune response to these released antigens, thereby improving the effectiveness of chemotherapy. The therapeutic activity of CAR-T cells has also been limited in most solid tumors, primarily due to poor *in vivo* persistence(34),(35). Accumulating evidence indicates that the co-expression of IL-15 enhances the proliferation, persistence, and anti-tumor capabilities of CAR-T cells. A recent clinical study showed that IL-15-armored GPC3 CAR-T cells significantly increased the proportion of memory and effector cells, leading to improved anti-tumor efficacy(36). Wu et al. also reported that a PD1-IL2Ra-IL2 can effectively augment the efficacy of CAR-T cells in solid tumors(18). In addition, our previous studies observed that tumor-conditional IL-15 can synergize with OT-I CD8^+^ T cells, resulting in the complete cure of MC38-OVA tumor in a mouse model(15). The combination therapy of PMIS and CAR-T cells holds tremendous promise for treating solid tumors.

Our study has limitations. Although we have observed that blocking CD25 enhances the antitumor activity of PSI but inhibits the antitumor efficacy of PMIS, indicating that pre-existing intratumoral CD25^+^CD8^+^ T cells play a crucial role in the effects of PD-1 cis-delivery cytokines, the optimal balance of CD25^+^CD8^+^ T cells to Tregs requires further studied. Additionally, PMIS can be cleaved within tumors to release free ILR, which can impact various immune cells beyond just T cells, necessitating comprehensive investigation. The effective activation of intratumoral CD25^+^CD8^+^ T cells depends on the enzymatic cleavage efficiency of PMIS. It is crucial to investigate whether the cleavage efficiency and the efficacy of PMIS vary among different tumor types.

In summary, our findings highlight the negative impact of Tregs on antitumor efficacy of cis-delivering cytokines and underscore the need for a higher CD25^+^CD8/Treg ratio to enhance antitumor immunity. The development of conditionally activated anti-PD-1/IL-15 presents potential solutions for improved efficacy as well as reduced toxicity. PMIS not only displays robust single-agent antitumor efficacy by revitalizing preexisting and exhausted CD8^+^ TILs but also can be effectively combined with other therapies to control orthotopic tumors and their metastases. The conditional cis-delivery of IL-15 superagonist overcomes nearly all limitations associated with current cytokines cis-delivery strategies, holding high translational promise for treating patients with cold and metastatic tumors.

## Materials and Methods

### Protein production

Anti-PD-L1/IL-15 (LH01) and non-targeting Fc-IL-15 (LH02) were constructed and produced as previously described(20). For unmasked anti-PD-1/IL-15 (PSI) construction, the human IL-15Rα-sushi domain (Ile31 to Val 115)/IL-15 mutant (IL-15N72D) complex (RLI) was directly fused to the C-terminal of anti-PD-1 heavy chain. For PIS or PMIS construction, the human IL-15 mutant (IL-15N72D)/ IL-15Rα-sushi domain (Ile31 to Val 115) complex (ILR) was fused to the C-terminal of anti-PD-1 heavy chain via a (GGGGS)_3_ (non-substrate) linker or a GSSGGSGGSGGSG-SGQLLGFLTA-GSSGGSGGSGGSG (substrate) linker, respectively. All plasmids were constructed by inserting DNA fragments into the previously used pMF09 vector(20). The plasmids were mixed with 25 kDa linear polyethylenimine and transiently transfected into HEK293E cells. All fusion proteins were purified using a protein A affinity column (GE Healthcare) and analyzed on SDS-PAGE under reducing conditions.

### Cell lines

HEK293E, CLTT-2, and Mo7e cell lines were kept in our laboratory and cultured as previous described(20). MC38, B16-F10, RM-1, and 4T1 cells were cultured in Dulbecco’s modified Eagle’s medium (DMEM) containing 10% FBS. All cells mentioned above were maintained under aseptic conditions and incubated at 37°C with 5% CO_2_.

### Animal experiments

All animal experiments were approved by the Animal Care and Use Committee of Shanghai Jiao Tong University (Ethics number: A2023153-004). Balb/c and C57BL/6 mice aged 6-8 weeks were purchased from Shanghai SLAC Laboratory Animal Co., Ltd. Male NCG mice aged 4-6 weeks were purchased from Jiangsu GemPharmatech LLC. All mice were raised in pathogen-free environments and received humane treatment throughout the experimental period. In the antitumor studies, tumors were measured every two or three days using a digital caliper, with volumes calculated as (length×width^2^)/2. The human endpoint for tumor size was 2000 mm^3^, except in cases of B16-F10 tumor models, which grew rapidly and could double in volume every other day.

### Establishment of PDX colon tumor model

Fresh colon cancer samples surgically resected were provided with approval from the Medical Research Ethics Committee of Hangzhou First People’s Hospital (No.KY-2024466-01). All subjects provided broad informed consent for the research use of their biological samples. Tumor tissue was washed with saline, cut into small pieces (approximately 2-3 mm), and transplanted subcutaneously into NCG mice. When tumors reach about 1 cm^3^, the mice were euthanized and the tumor was excised. Tumor pieces were then harvested and re-implanted into new NCG mice to create a secondary PDX model. Mice in the fourth-generation, when the tumors grew to approximately 100 mm^3^, were used for drug efficacy studies.

### ELISA assessment of the affinity of anti-PD-1, PSI, or PMIS for PD-1

ELSIAs were conducted following standard procedures. Briefly, 96-well ELISA plates were coated overnight at 4 °C with 1.5 μg/mL of recombinant human or mouse PD-1 (Novoprotein, cat: CX91; cat: CJ98). The plates were then washed four times with PBST and blocked with 5% bovine serum albumin for 2 h at room temperature. After washing the plates, serial dilutions (1:3) of anti-PD-1, PSI, or PMIS were added in duplicate to the plates and incubated at room temperature for 2 h. The plates were then washed four times and incubated with Peroxidase AffiniPure Goat Anti-Human IgG (H+L) (1:10000 dilution) for 1 h at room temperature. Following another wash, the plates were incubated in the dark with TMB single-component substrate solution for 3-5 min. The reaction was stopped with 2 M sulfuric acid, and absorbance was read at 450 nm with a reference at 630 nm (Teacan, Infinite 200 PRO).

### *In vitro* cleavage of PMIS with MMP-2

Human MMP-2 (SinoBiological, cat: 10082-HNAH) was firstly activated via incubation 1 μg of protease with 0.2 mM APMA at 37°C for 1 h in TCNB activation buffer (50 mM Tris, 10 mM CaCl2, 150 mM NaCl, 0.05% (w/v) Brij 35, pH7.5). Subsequently, 25 μg PMIS was incubated with the reaction buffer at 37°C for 12 h.

### Detection of plasma ALT, AST, and IFN-γ

Plasma levels of ALT and AST were measured using a Roche biochemical analyzer (Roche, Switzerland). Plasma level of IFN-γ was determined by mouse IFN-γ ELISA Kit (PeproTech, cat: 900-T98K) according to the manufacturer’s procedure.

### Quantitative biodistribution studies of immunocytokine

Heart, liver, spleen, lung, kidney, and tumor tissues from B16-F10 tumor-bearing mice were collected and minced. About 100-150 mg of each tissue was weighed and homogenized in 10% PBS before being centrifuged to obtain the supernatant. The above developed ELISA assay for measuring anti-PD-1/IL-15 affinity for PD-1 was used to detect the total amount of PSI or PMIS in each homogenate.

### Fluorescence imaging

PMIS was labeled with SulfoCy5.5 SE (Fluorescence, cat: 1057-1 mg) and excess dye unbound PMIS was removed via ultrafiltration. Fluorescently labeled PMIS (1 mg/kg) was intravenously injected into tumor-bearing mice. Fluorescence was measured with PE IVIS Spectrum at different time points.

### Flow cytometry analysis

About 200 mg of tumor tissues was minced and resuspended in digestion buffer [RPMI1640 medium containing collagenase IV (2 mg/mL) (Yeasen, cat: 40510ES76) containing hyaluronidase (1.2 mg/mL) (Yeasen, cat: 20426ES80)]. Tumors were digested for 60 min at 37°C and then filtered through a 200-mesh nylon net to obtain the cell suspension. The cells were washed with RPMI 1640, filtered through a 200-mesh nylon net again, and then resuspended in PBS to obtain pre-treated single cell suspension. Splenic lymphocytes were isolated from the spleens with lymphocyte separation medium (DAKEWE, cat: 7211011) after the spleens were gently ground.

The Zombie Aqua Fixable Viability Kit was used to exclude dead cells and was incubated with cell samples at 25°C for 15 min. Samples were then blocked with anti-mouse CD16/CD32 mAb 2.4G2 at 4°C for 30 min before being incubated with surface marker antibodies at 4°C for an additional 30 min. For the detection of intracellular markers, cell samples were further fixed and permeabilized usng the Transcription Factor Buffer Set. Flow cytometry was performed using CytoFLEX cytometer (Beckman Coulter, USA), ACEA Novocyte (Agilent, Technologies, USA), or Cytek Aurora (Cytek Biosciences, USA) and analyzed with FlowJo 10.6.1 (TreeStar, USA). Gate margins were establishd using isotype controls and fluorescence-minus-one controls. Antibodies and reagents for flow cytometry are shown in Table S1.

### Depletion of immune cells in mice

In detail, to deplete the individual immune cell types, mice were intraperitoneally given 200 μg of anti-NK1.1 antibody (BioXCell, Cat: BE0036), 200 μg of anti-CD8α antibody (BioXCell, Cat: BE0061), , or 100 μg of anti-CD25 antibody (BioXcell, Cat: BE0012). To study the effect of lymphocytes egress from lymph nodes, FTY720 (Sigma-Aldrich, Cat: 162359-56-0) was administered intraperitoneally at a dosage of 1.25 mg/kg.

### RNA sequencing

Total RNA was extracted from B16-F10 tumor tissues. cDNA library construction, sequencing, and data analysis were performed by Shanghai Majorbio Bio-Pharm Technology Co., Ltd. using the Majorbio cloud platform. High-quality reads were aligned to the mouse reference genome (GRCm39) using Bowtie2. Gene expression levels were normalized to fragments per kilobase of exon model per million mapped reads (FPKM) based on the expectation-maximization method. The NOISeq method was used to screen out differentially expressed genes (DEG), with statistical significance defined as a fold change > 1.5 and p values < 0.05. We accessed the Gene Set Enrichment Analysis website to obtain gene sets associated with immunity. Immune signature scores were calculated as the mean log_2_ (fold change) across all genes in each signature. The FPKM values of these genes were logarithmically (fold-change) converted, and heat maps were generated using GraphPad Prism V.8 software.

### Preparation of mouse splenic T cells

Spleens from Balb/c mice were mechanically disrupted and gently ground through a 200-mesh nylon net again. The lymphocytes were isolated with lymphocyte separation medium (Dakewe, cat: DKW33-R0100). Add 1 μg/mL anti-mouse CD3 antibody (MultiSciences, cat: F2100300-100) and 2 μg/mL anti-mouse CD28 antibody (MultiSciences, cat: F2102800-100) to a 24-well plate and coat overnight at 4°C. Wash the plate with PBS to remove unbound antibodies. Splenocytes were then resuspended at a cell density of 1.5 × 10^6^/mL in complete RPMI medium supplemented with mouse IL-2 (10 ng/mL) and 2-mercaptoethanol (50 μM). Activated T cells were obtained after a 72 h culture. Adjust the cell density to 1.0 × 10^6^/mL, and plate 2 × 10^5^ cells in a 48-well plate. The cells were treated with PSI (0.02, or 0.2 μg/mL), or cleaved PMIS (1.0, or 2.0 μg/mL). After 48 hours, proceed with flow cytometry analysis.

### Immunofluorescence analysis

Tumor tissues were fixed in 4% paraformaldehyde, embedded in paraffin, and sectioned (4 μm). Co-staining of CD8 and Foxp3 in the B16-F10 tumor sections was performed using a Three color mIHC Fluorescence kit (Recordbio, cat: RC0086-23) based on tyramide signal amplification (TSA) technology according to the manufacturer’s instruction. Sections were imaged using a Pannoramic DESK slide scanner.

### Statistical analysis

Prism 8.0 software (GraphPad, USA) was used for statistical analysis. The unpaired two-tailed Student’s t test and one-way ANOVA were used to determine the statistical significance of differences between experimental groups (*: *p* < 0.05, **: *p* < 0.01, ***: *p* < 0.001). The log rank (Mantel-Cox) test was used to assess survival.

## Data Availability Statement

The datasets used or analyzed in this study are available from the corresponding author upon request.

## Acknowledgements

The work was partly supported by the National Natural Science Foundation of China (82404500 to Dr. Shi W, W2412118 to Dr. Lu H, and 82203708 to Dr. Kong W), the Natural Science Foundation of Chongqing (2022NSCQ-MSX2319 to Dr. Lu H), the Science & Technology Commission of Shanghai Municipality (No. 23ZR1431800 to Dr. Lu H), and Zhejiang Medical and Health Technology Project (2023RC058, 2024KY175 to Dr. Kong W).

## Author Contributions

**Q. Zeng:** Conceptualization, Writing review, formal analysis, investigation, methodology. **W. Kong:** Funding acquisition, resources. **Y. Qin:** Investigation. **L. Song:** Investigation, methodology. **N. Liu:** Writing-review and editing, discussion. **Z Liu:** Investigation, methodology. **C. Ji:** Investigation, methodology. **G. Qi:** Investigation, validation. **W. Shi:** Conceptualization, methodology, writing-original draft, writing-review and editing, funding acquisition. **H. Lu:** Conceptualization, resources, funding acquisition, supervision, writing-review and editing.

## Declaration of Interests

The authors declare no competing interests.

## Notes

### Competing Interest Statement

The authors have declared no competing interest.

