## Supplemental Figures 1-7 and Table 1 for "Cis-delivering releasable IL-15 superagonist enhances antitumor immunity in cold tumors by invigorating preexisting CD25^+^CD8^+^ T cells"

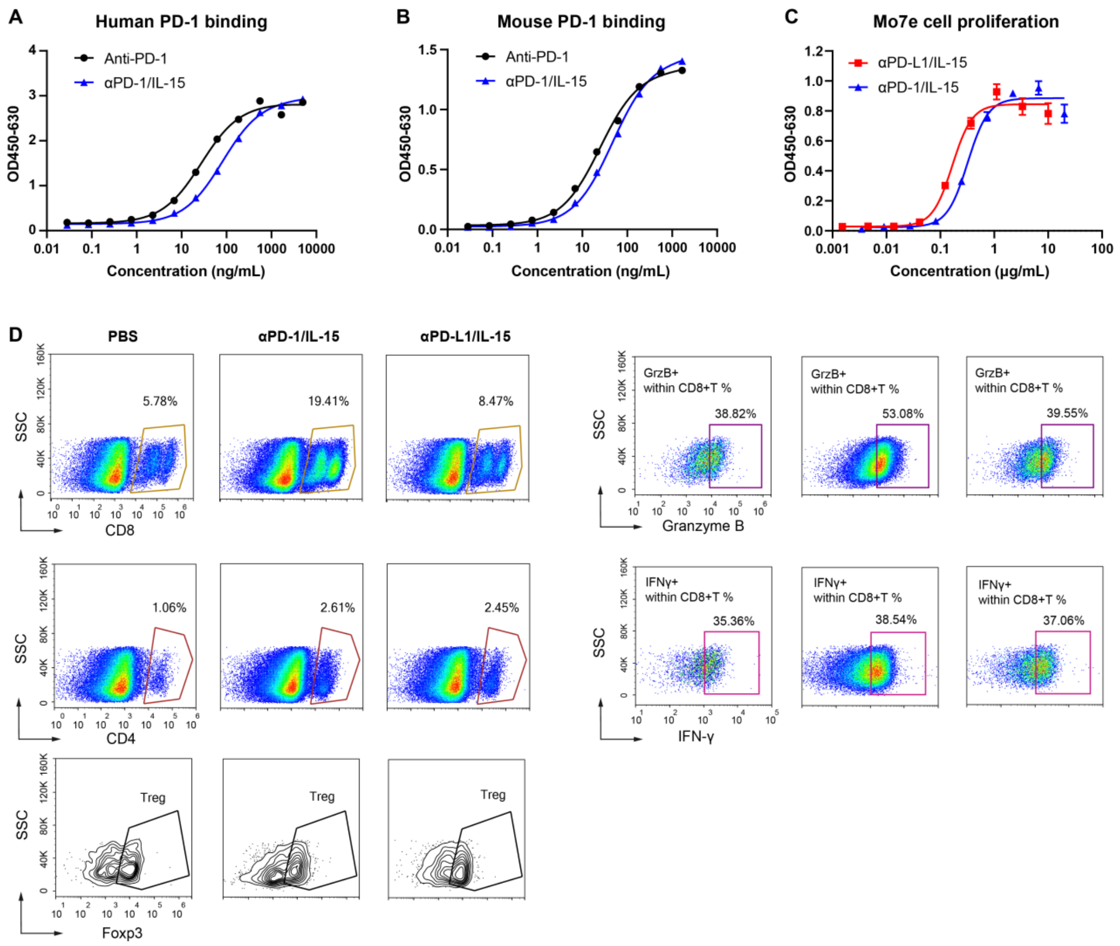


**Figure S1. Cis-delivery showed superior antitumor effects to trans presentation but with higher toxicity, related to Figure 1.**

(A and B) Binding of anti-PD-1 and αPD-1/IL-15 to plate-bound human or mouse PD-1 (n = 2 technical replicates). Data were analyzed using the one site-total to calculate the $EC_{50}$ values.

(C) The proliferative potential of αPD-L1/IL-15 and αPD-1/IL-15 in human Mo7e cells (n = 3 technical replicates). Data were analyzed using the four-parameter-fit logistic equation to calculate the $EC_{50}$ values. The data is shown as mean ± SEM.

(D) Flow cytometry analysis of dissociated tumors.

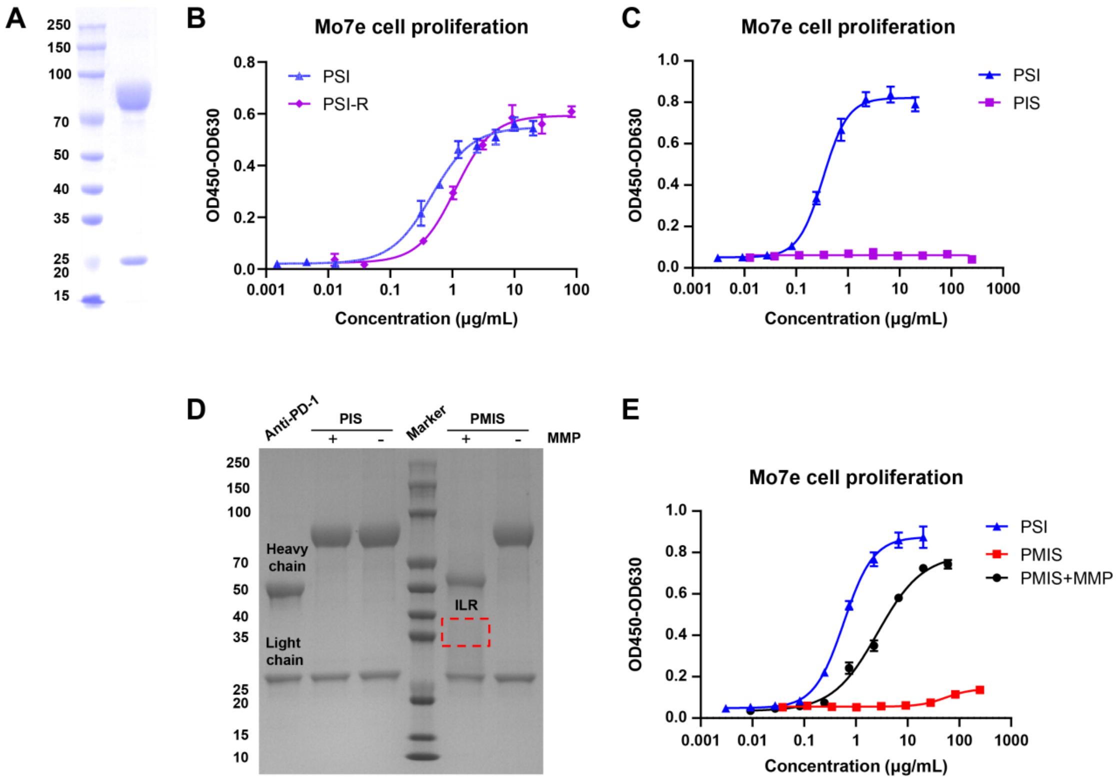


**Figure S2. Engineering and characterization of a PD-1-targeted and conditionally-activated IL-15 superagonist**

(A) Reducing SDS-PAGE analysis of PSI-R.

(B and C) The proliferative potential of PSI, PSI-R (B), and PIS (C) in human Mo7e cells (n = 3 technical replicates). Data were analyzed using the four-parameter-fit logistic equation to calculate the $EC_{50}$ values. The data is presented as the mean ± SD.

(D) Reducing SDS-PAGE analysis of anti-PD-1, PIS or PMIS as well as MMP2-cleaved PIS or PMIS. The experiment was repeated three times independently with similar results.

(E) The proliferative potential of PSI, PMIS, and PMIS + MMP in human Mo7e cells (n = 3 technical replicates). Data were analyzed using the four-parameter-fit logistic equation to calculate the $EC_{50}$ values. The data is presented as the mean ± SD.

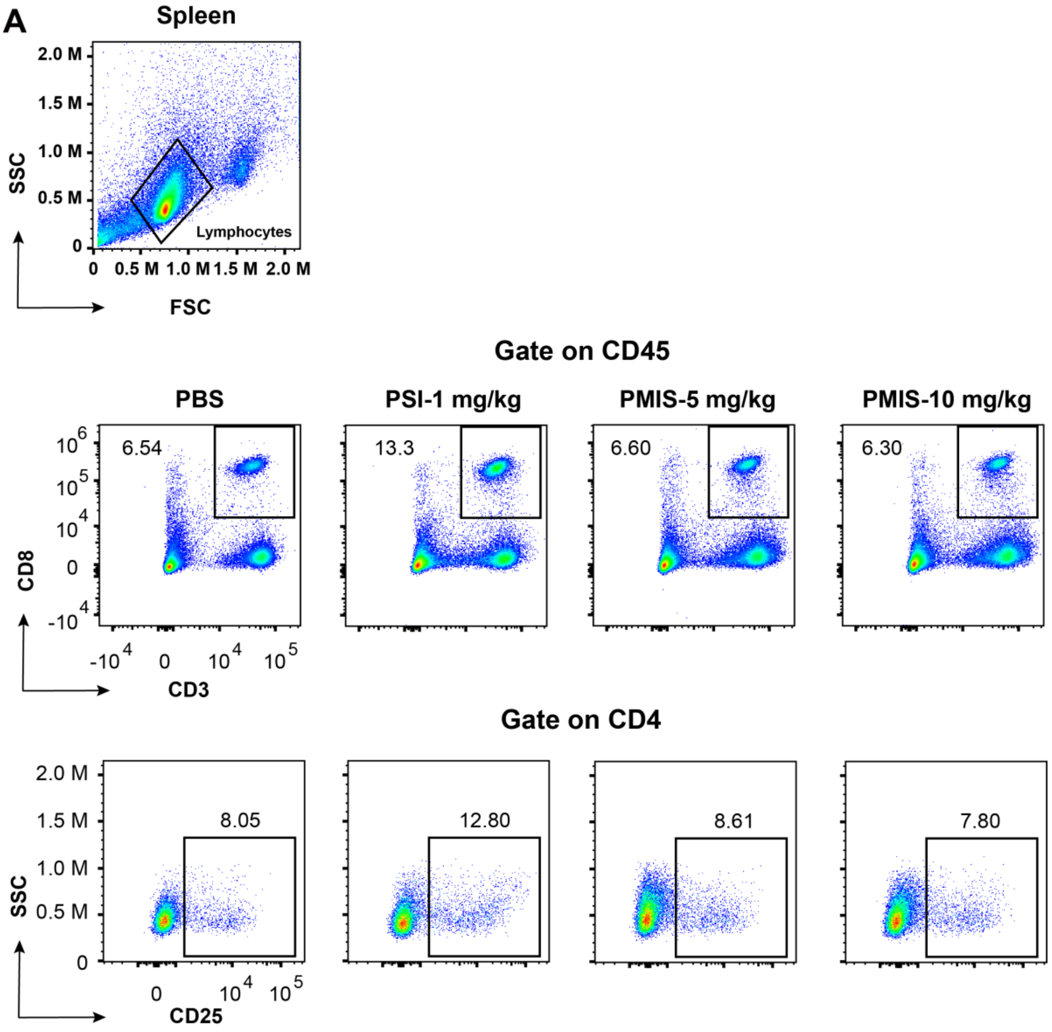


**Figure S3. PMIS significantly reduces systemic toxicity associated with IL-15, related to Figure 3.**

(A) The gate to identify lymphocytes of spleen. The percentages of splenic $CD8^+$ T cells and $CD4^+CD25^+$ T cells for $CD45^+$ or $CD4^+$ lymphocytes were shown, respectively.

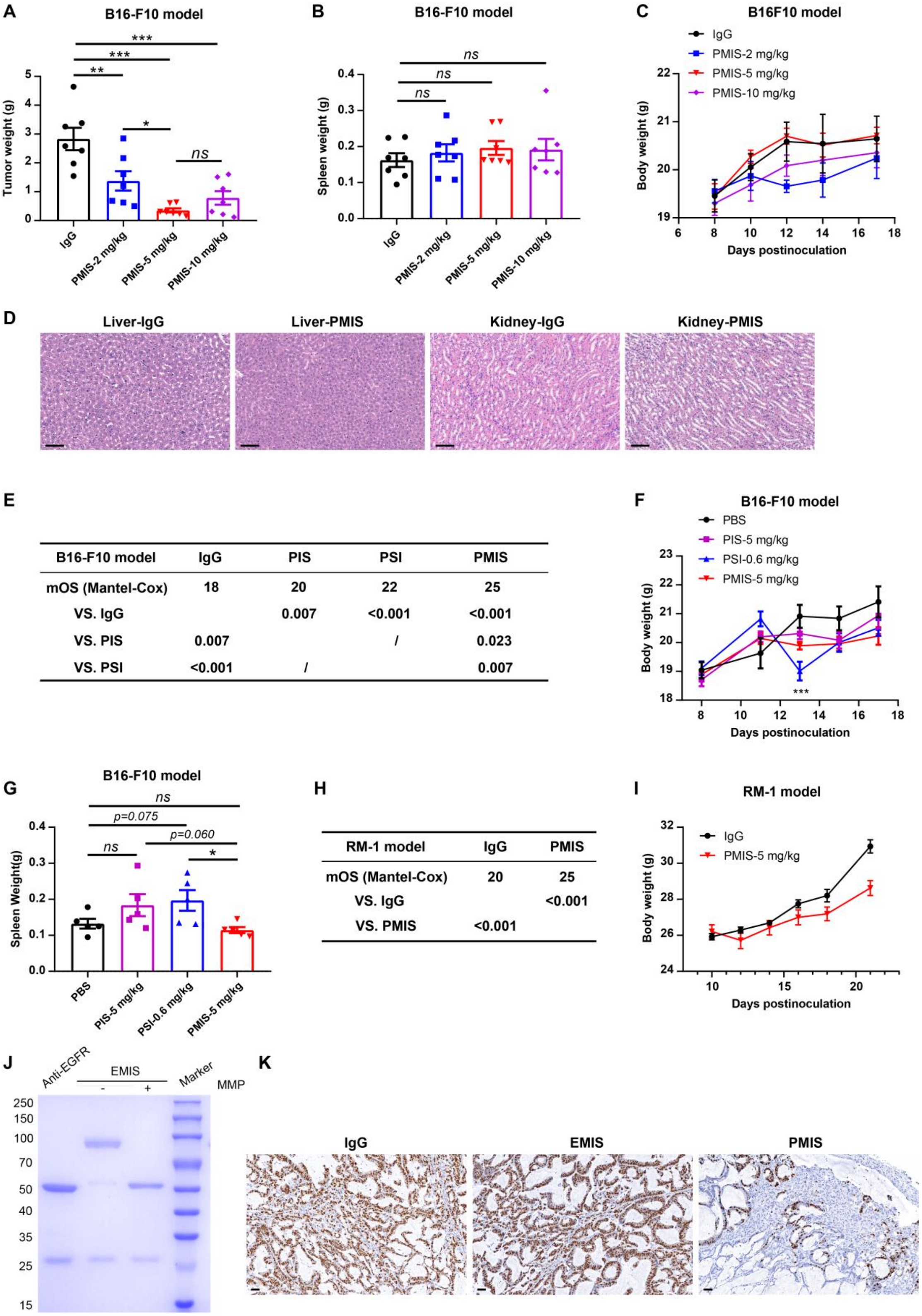


| B16-F10 model | IgG | PIS | PSI | PMIS |
|---|---|---|---|---|
| mOS (Mantel-Cox) | 18 | 20 | 22 | 25 |
| VS. IgG | | 0.007 | <0.001 | <0.001 |
| VS. PIS | 0.007 | | / | 0.023 |
| VS. PSI | <0.001 | / | | 0.007 |

| RM-1 model | IgG | PMIS |
|---|---|---|
| mOS (Mantel-Cox) | 20 | 25 |
| VS. IgG | | <0.001 |
| VS. PMIS | <0.001 | |

**Figure S4. PMIS enhances antitumor effects with significantly improved safety, related to Figure 4.**

(A-D) B16-F10 tumor cells ($4 \times 10^5$) were subcutaneously implanted into female

C57BL/6 mice. Mice were then randomized into four groups, and treatment initiated when tumors reached 50-100 $mm^3$. On days 8, 11, and 14 (n = 7), mice were intravenously injected with IgG control (5 mg/kg) or PMIS (2 mg/kg, 5 mg/kg, or 10 mg/kg). On day 17, the mice were euthanized, and both tumors and spleens were excised and weighed (A and B). Body weight of tumor-bearing mice was monitored (C). H&E staining was performed for livers and kidneys of IgG or PMIS group (10 mg/kg) (scale bar: 50 μm) (D).

(E-G) B16-F10 tumor cells ($4\times10^5$) were subcutaneously implanted into female C57BL/6 mice. Mice were then randomized into four groups, and treatment initiated when tumors reached 50-100 $mm^3$. On days 8, 11, and 14 (n = 9-10), mice were intravenously injected with PBS control, PIS (5 mg/kg), PSI (0.6 mg/kg), or PMIS (5 mg/kg). Survivals were plotted and analyzed (E). The body weights of tumor-bearing mice were recorded (F). The spleens of mice (n = 5) were extracted and weighed after euthanasia (G).

(H and I) RM-1 tumor cells ($5\times10^5$) were subcutaneously implanted into male C57BL/6 mice. Mice were then randomized into two groups, and treatment initiated when tumors reached 50-100 $mm^3$. On days 10, 13, and 16 (n = 8), mice were intravenously injected with IgG control or PMIS (5 mg/kg). Survival was shown and analyzed (H). The body weights of mice were monitored (I).

(J) Reducing SDS-PAGE analysis of anti-EGFR, or EMIS as well as MMP2-cleaved EMIS. The experiment was repeated three times independently with similar results.

(K) Immunohistochemical staining for Ki67 was performed on the tumor tissues of the PDX tumor model (scale bar: 50 μm).

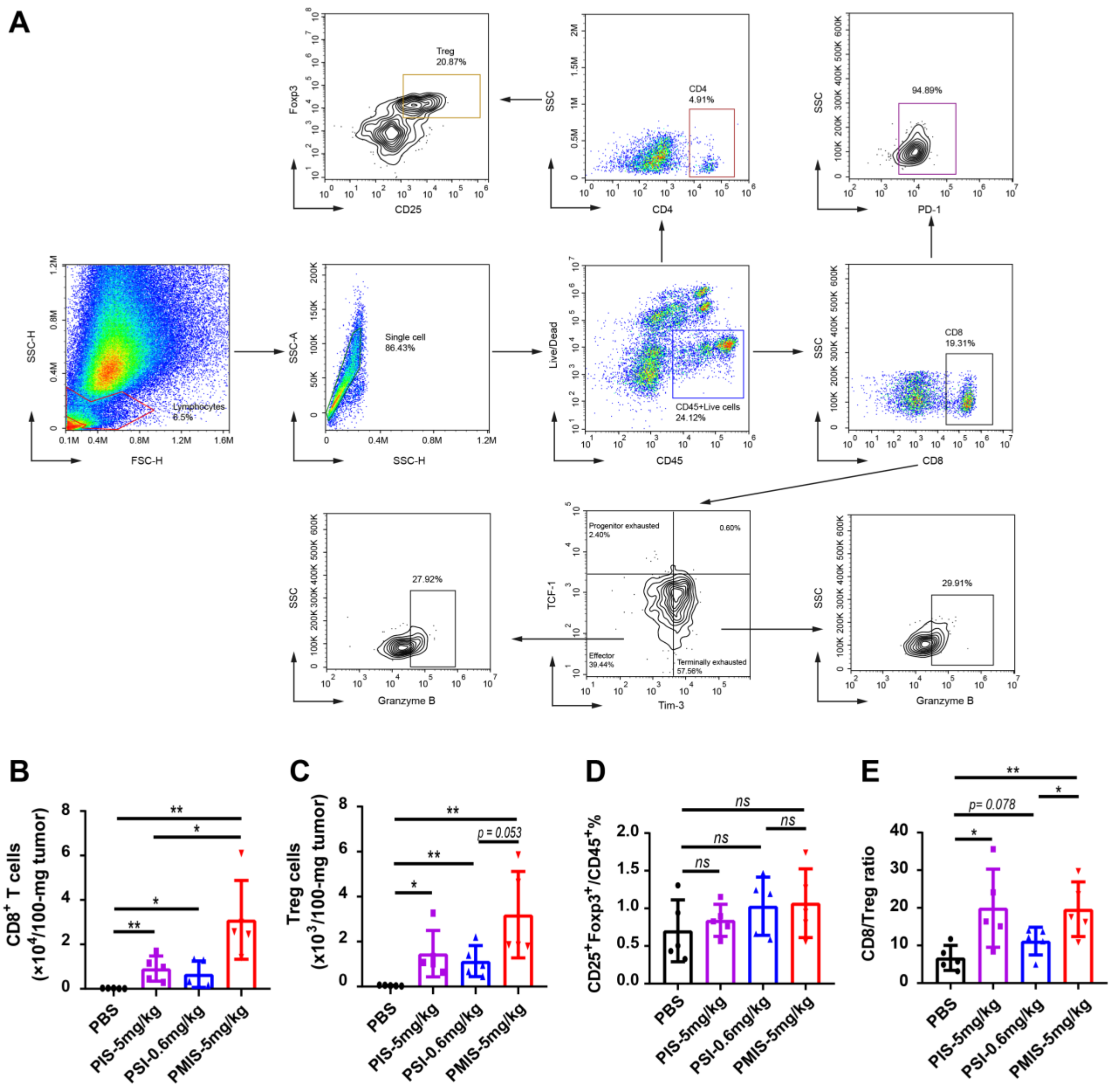


**Figure S5. PMIS markedly increases CD25$^{+}$CD8/Treg ratio within tumor, related to Figure 5.**

(A) Representative gating strategy for identifying CD8$^{+}$ T cells, Tregs, progenitor exhausted CD8$^{+}$ T cells, effector CD8$^{+}$ T cells, terminally exhausted CD8$^{+}$ T cells, granzyme B$^{+}$ effector CD8$^{+}$ T cells, and granzyme B$^{+}$ terminally exhausted CD8$^{+}$ T cells in the B16-F10 tumor tissue.

(B and C) The numbers of intratumoral CD8$^{+}$ T cells (B) and Tregs (C) were shown (n = 5).

(D) The percentage of intratumoral Tregs was shown for populations of CD45$^{+}$

lymphocytes (n = 5).

(E) CD8/Treg ratio is calculated.

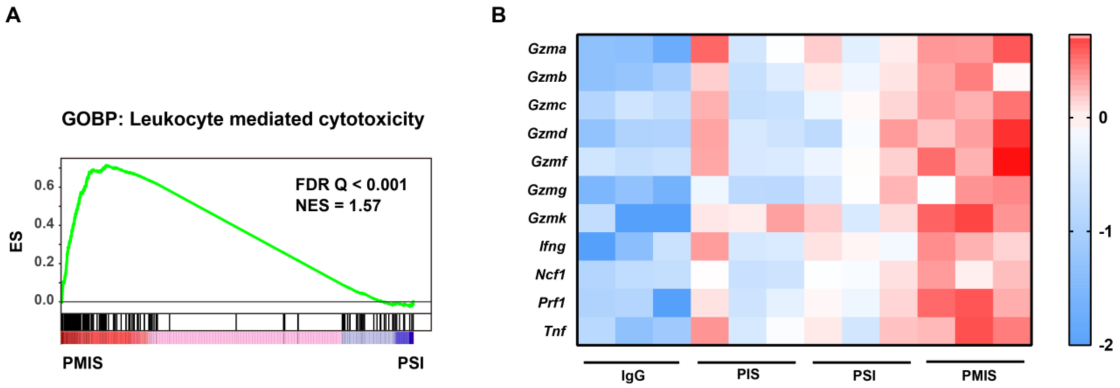


**Figure S6. The antitumor efficacy of PMIS depends on intratumoral CD25$^+$CD8$^+$ T cells, related to Figure 6.**

(A) Gene set enrichment analysis of leukocyte-mediated cytotoxicity after PMIS or PSI treatment.

(B) The heatmap depicts gene expression alterations of leukocyte-mediated cytotoxic effectors in response to different treatments.

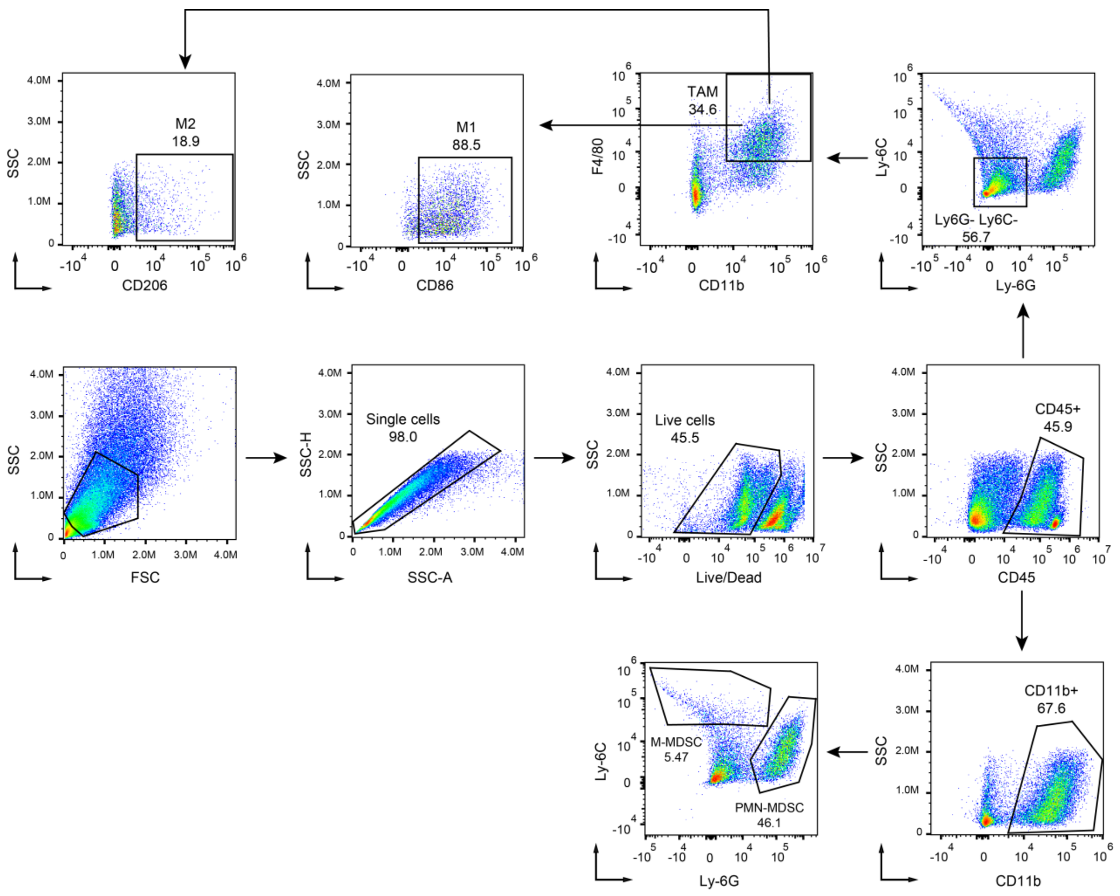


**Figure S7. PMIS can significantly inhibit orthotopic cold carcinoma and its metastasis, related to Figure 7.**

Representative gating strategy for identifying PMN-MDSCs, M-MDSCs, M1 macrophages, and M2 macrophages in the 4T1 tumor tissue.

**Table S1.** Reagents and antibodies for flow cytometry analysis

| **Reagents** | **Source** | **Identifier** |
|---|---|---|
| APC/Cy7 anti-mouse CD45.2 (clone:104 ) | BioLegend | Cat#: 109824; RRID: AB_830789 |
| FITC anti-mouse CD3ε (clone:500A2) | BioLegend | Cat#: 152304; RRID: AB_2632667 |
| APC anti-mouse CD8a (clone: 53-6.7) | BD Biosciences | Cat#: 553035; RRID: AB_398527 |
| PE anti-mouse TIM-3 (clone: 25F.1D6) | BD Biosciences | Cat#:568904 |
| PE/Cy7 anti-human/mouse Granzyme B (clone: QA16A02) | BioLegend | Cat#: 372213; RRID: AB_2728380 |
| Alexa Flour 647 anti-mouse TCF-7/TCF-1 (clone: S33-966) | BD Biosciences | Cat#: 566693; RRID: AB_2869823 |
| Fc block-anti-mouse CD16/32 | BioLegend | Cat#: 101302; RRID: AB_312801 |
| Zombie Red Fixable Viability Kit | BioLegend | Cat#: 423109 |
| Transcription Factor Buffer Set | BD Biosciences | Cat#: 562574; RRID: AB_2869424 |
| PE anti-mouse CD4 (clone:GK1.5) | BD Biosciences | Cat#:557308; RRID:AB_396634 |
| PE-CF594 anti-mouse CD8a (clone:53-6.7) | BD Biosciences | Cat# 562283; RRID:AB_11152075 |
| BV421 anti-mouse CD25 (clone: PC61) | BioLegend | Cat# 102033; RRID:AB_10895908 |
| Alexa Flour 647 anti-mouse Foxp3(clone: 150D) | BioLegend | Cat# 320013; RRID:AB_439749 |
| BV786 anti-mouse PD-1 (clone:J43) | BD Biosciences | Cat# 744548; RRID:AB_2742319 |
| BV650 anti-mouse IFN-γ (clone:XMG1.2) | BD Biosciences | Cat# 563854; RRID:AB_2738451 |
| BV421 anti-mouse CD206 (clone: C068C2) | BioLegend | Cat# 141714; RRID:AB_10917384 |
| PE anti-mouse CD86 (clone:GL-1) | BioLegend | Cat# 105007; RRID： AB_313150 |
| FITC anti-mouse CD11b (clone:M1/70) | BioLegend | Cat# 101205; RRID:AB_312788 |
| PerCP/Cyanine5.5 anti-mouse Ly-6C (clone:HK1.4) | BioLegend | Cat# 128011; RRID:AB_1659242 |
| APC anti-mouse Ly6G (clone: 1A8) | BioLegend | Cat# 127613; RRID:AB_1877163 |

| | | |
|---|---|---|
| PE/Cyanine7 anti-mouse F4/80 (clone: BM8) | BioLegend | Cat# 123113; RRID:AB_893490 |
